# Astrocytic connexin43 phosphorylation contributes to seizure susceptibility after mild traumatic brain injury

**DOI:** 10.1101/2024.11.12.623104

**Authors:** Carmen Muñoz-Ballester, Owen Leitzel, Samantha Golf, Chelsea M Phillips, Michael J Zeitz, Rahul Pandit, Elizabeth Wash, Jenna V. Donohue, James W. Smyth, Samy Lamouille, Stefanie Robel

## Abstract

Astrocytes play a crucial role in maintaining brain homeostasis through functional gap junctions (GJs) primarily formed by connexin43 (Cx43) in the cortical gray matter. These GJs facilitate electrical and metabolic coupling between astrocytes, allowing the passage of ions, glucose, and metabolites. Dysregulation of Cx43 has been implicated in various pathologies, including traumatic brain injury (TBI) and acquired epilepsy. After mild TBI/concussion, we previously identified a subset of atypical astrocytes, which are correlated with the development of spontaneous seizures. These astrocytes exhibit reduced Cx43 expression and coupling. However, atypical astrocytes represent a relatively small subset of astrocytes within the cortical gray matter and previous studies suggest an overall increase of Cx43 protein after TBI. Additionally, Cx43 also has non-junctional and channel-independent functions, which include hemichannel communication with the extracellular milieu, cell adhesion, protein trafficking, protein-protein interactions, and intracellular signaling. In the present study, we set out to determine how mild TBI initiates alterations to Cx43 protein expression and localization, how they may be regulated, and whether they contribute to seizure susceptibility. We demonstrate remarkable heterogeneity of Cx43 protein levels from astrocyte to astrocyte. In accordance with our previous findings, a subset of astrocytes lost Cx43 expression, yet total cortical Cx43 protein increased. At the subcellular level, junctional Cx43 protein levels remained stable, while hemichannels and/or cytoplasmic Cx43 were increased. Phosphorylation of Cx43 at serine 368, a key regulatory site influencing GJ assembly and function, increased after mild TBI. Critically, Cx43^S368A^ mutant mice, lacking this phosphorylation, exhibited reduced susceptibility to pentylenetetrazol-induced seizures. These findings suggest that TBI-induced Cx43 phosphorylation enhances seizure susceptibility, while inhibiting this modification presents a potential therapeutic avenue for mitigating neuronal hyperexcitability and seizure development.

**Significance statement:** Connexin43 (Cx43) is the main protein comprising astrocyte gap junctions which mediate astrocyte coupling into cellular networks, but it also has other non-junctional functions. Many pathologies present with altered Cx43 regulation. In this study, we assessed Cx43 alterations after mild traumatic brain injury (TBI) in a mouse model. We found that while some astrocytes lost Cx43 expression, other astrocytes had increased cytoplasmic and hemichannel Cx43. This increase correlated with an increase in phosphorylated Cx43 at serine 368. Cx43^S368A^ mutant mice, lacking this phosphorylation, exhibited reduced susceptibility to seizures induced by pentylenetetrazol (PTZ). These findings suggest that TBI-induced Cx43 phosphorylation enhances seizure susceptibility.

## Introduction

Astrocytes maintain brain homeostasis in part through functional gap junctions (GJs), comprised of connexin43 (Cx43), which mediate the coupling of astrocytes into vast cellular networks (Rouach et al., 2008; Visser et al., 2024; Wallraff et al., 2006; Xu et al., 2010). Cx43 is one of two primary GJ proteins expressed in astrocytes in the brain (Magnotti et al., 2011) with connexin30 (Cx30) being expressed at significantly lower levels in cortical gray matter astrocytes (Rash et al., 2001). GJ coupling is electrical and metabolic, since ions, glucose, and other metabolites can pass from the cytoplasm of one astrocyte to the next. Many pathologies (Gajardo-Gómez et al., 2017; Masaki, 2015; Orellana et al., 2011; Sarrouilhe et al., 2017; Vis et al., 1998; Wang et al., 2018), including traumatic brain injury (TBI) (B. Chen et al., 2017; W. Chen et al., 2019; Wu et al., 2013; Xia et al., 2024) and acquired epilepsy (Bedner et al., 2015; Deshpande et al., 2017; Walrave et al., 2018) present with altered Cx43 regulation, and are often equated with GJ dysfunction. Yet, Cx43 also has non-junctional and channel-independent functions, which include hemichannel communication with the extracellular milieu, cell adhesion, protein trafficking, protein-protein interactions, and intracellular signaling (Martins-Marques et al., 2019).

We recently reported that Cx43 protein expression, as well as astrocyte coupling, decreases in an atypical subset of astrocytes after mild TBI, and these atypical astrocytes are associated with the development of spontaneous seizures (Shandra et al., 2019). TBI affects between 64 and 69 million people worldwide each year (Dewan et al., 2018). Approximately 70% of TBI cases are mild, and yet, even mild TBI can lead to long-term consequences, such as impaired cognitive function, emotional regulation, and sleep (Mantua et al., 2017). Mild TBI also increases the risk for developing co-morbidities, including post-traumatic epilepsy (Sødal et al., 2024; Verellen & Cavazos, 2010). Currently, no clinical interventions are available that prevent those consequences after TBI. This is in part because the early molecular and cellular events initiated by mild TBI that lead to long-term consequences have not been fully resolved.

Reduced Cx43-mediated astrocyte coupling is thought to contribute to neuronal dysfunction, potentially through impaired spatial potassium (K^+^) buffering. When K^+^ is elevated in the extracellular space, neurons depolarize, which can result in hyperexcitability (David et al., 2009; Feng & Durand, 2006; Steinhäuser et al., 2012; Ullah et al., 2009). When the delicate balance of neuronal inhibition and excitation is disrupted, lack of Cx43/Cx30 appears to promote neuronal hyperexcitability and seizures in a mouse model of epilepsy induced by kainate injection (Deshpande et al., 2020) and the seizure-inducing drug pentylenetetrazol (PTZ) causes more frequent seizures (Chever et al., 2016a). In this latter study, seizures were milder and shorter, possibly due to the role of astrocyte GJ in distributing glucose and lactate to promote and sustain neuronal activity (Rouach et al., 2008).

Many studies assessing the role of astrocyte Cx43 in pathology were conducted in mice with widespread deficiency in both Cx30 and Cx43 (typically Cx30 full knockout (KO) and Cx43 conditional KO using human-GFAP-Cre, which is expressed during mid-embryonic development in radial glia) (Chever et al., 2016b; Deshpande et al., 2020; Rouach et al., 2008). Yet, after mild TBI, our previous study showed reduction of Cx43 only in a small subset of atypical astrocytes while Cx43 was still present in most astrocytes even in those mice that developed seizures (Shandra et al., 2019). As most previous studies that assessed Cx43 used bulk analysis, this heterogeneity has remained unexamined in the context of TBI.

Further, conditional Cx43 KO mice (Chever et al., 2016b; Deshpande et al., 2020; Rouach et al., 2008) and Cx43 siRNA (Ichkova et al., 2019) both target Cx43 transcripts, and thus affect Cx43 GJ channel-dependent and -independent functions. Cx43 can also remain undocked, forming hemichannels which may release small molecules, such as glutamate or ATP into the extracellular space driving pathology (Orellana et al., 2011; Retamal et al., 2006; Stout et al., 2002; Ye et al., 2003). Recent studies demonstrate that astrocytic Cx43 hemichannels increase hyperexcitability and neuronal damage in the context of amyotrophic lateral sclerosis (ALS) (Almad et al., 2022) and temporal lobe epilepsy (Guo et al., 2022).

GJs are dynamic structures that turn-over constantly under normal conditions and therefore remodel rapidly after insult due to alterations to rates of turnover and/or synthesis and forward trafficking of *de novo* connexin protein (Hervé et al., 2007). Cx43 localization and activity are regulated by several kinases modulating the phosphorylation of key residues within the Cx43 carboxy terminus (Lampe & Laird, 2022). Phosphorylation of Cx43 at serine 368 (pCx43^S368^) is of particular interest for epilepsy and TBI because pCx43^S368^ is increased in animal models and patients (W. Chen et al., 2018; Deshpande et al., 2017; Greer et al., 2017). Of the known residues subject to dynamic phosphorylation in the Cx43 C-terminus, pCx43^S368^ is well accepted as a key indicator of alterations to gap junction function and stability in response to stressors. Not only is it associated with reduced channel opening probability, but it is a key component of a phosphorylation ‘cascade’, which can elicit internalization and degradation of gap junctions (Cone et al., 2014; Smyth et al., 2014; Thévenin et al., 2013). Yet, despite correlative evidence suggesting a role for pCx43^S368^ in TBI and epilepsy, it remains unresolved whether pCx43^S368^ is causally linked to seizure susceptibility after TBI.

Cx43^S368^ phosphorylation is mediated by protein kinase C (PKC) and has been implicated in preventing new GJs from assembling, while negatively impacting channel opening probability (Calhoun et al., 2020; Ek-Vitorin et al., 2006; Lampe et al., 2000; Sirnes et al., 2009). During acute cardiac ischemia, pCx43^S368^ precedes ubiquitination of Cx43, leading to its degradation (Martins-Marques et al., 2015; Smyth et al., 2014). Mice harboring a serine to alanine mutation at Cx43^S368^ (Cx43^S368A^) to prevent phosphorylation have demonstrated protection against pathological cardiac conduction slowing during acute infection, attributed to maintenance of gap junction coupling (Gy et al., 2011; Padget et al., 2024).

Here we first assessed the heterogeneity of Cx43 protein levels after mild TBI. As previously reported, we confirmed a small subset of astrocytes had lost Cx43 protein expression (Shandra et al., 2019). Yet surprisingly, we observed an overall increase in Cx43 expression at three days after mild TBI with remarkable heterogeneity between groups of astrocytes. We determined that junctional Cx43 remains largely unchanged while the soluble fraction, which contains hemichannels and cytoplasmic Cx43, accounts for the increase in protein expression. Hemichannel activity was increased in the majority of but not all mice. Finally, we found increased pCx43^S368^ after mild TBI, and Cx43^S368A^ mice, which lack phosphorylation at Cx43^S368^, were less susceptible to seizures in response to PTZ kindling after mild TBI. Together, these results suggest that mild TBI induces pCx43^S368^, which confers a higher seizure susceptibility, and inhibition of pCx43^S368^ may have therapeutic potential in suppressing neuronal hyperexcitability and seizures following TBI.

## Material and methods

### Animals

Male and female mice were used for experiments. C57Bl/6 mice were bred in-house, and breeders were purchased from The Jackson Laboratory (JAX #000664). Cx43^S368A^ mice were kindly provided by Dr. Paul Lampe at the Fred Hutchinson Cancer Research Center (Freitas-Andrade et al., 2019). Aldh1l1-tdTomato astrocyte reporter mice (Tg(Aldh1l1-tdTomato)TH6Gsat) were bred in-house. Colonies were maintained in a standard pathogen restricted barrier animal facility in groups of 5 or 7 mice at maximum at the Fralin Biomedical Research Institute (Virginia Tech) or the University of Alabama at Birmingham, respectively. Animal facilities operated on a 12 h light/12 h dark cycle. Humidity and temperature were constant (22°C), with food and water provided ad libitum. After all procedures, mice were housed alone or with littermates that were a part of the same experimental condition until the desired endpoint was reached.

All animal procedures were approved and conducted according to the guidelines of the Institutional Animal Care and Use Committee of Virginia Polytechnic and State University and/or the University of Alabama at Birmingham. The protocols complied with the National Institute of Health’s Guide for the Care and Use of Laboratory Animals.

### Cx43^S368A^ mouse model

To determine the genotype of Cx43^S368A^ mice, EWLPF Primer (CTCACAGCTCTAATCCTCCTCCCTC) and EWLPR Primer (CTGACAGGCTCTACCTACCAAGTGC) (**Suppl. Fig. 1A**) were added at a final concentration of 1 µM in a solution with GoTaq G2 Green Master mix at 1x. Using the program described in **Suppl. Fig. 1B**, a band of 510 basepairs (bp) was visible for mutant animals while a band of 460 bp was visible in C57Bl/6 animals (**Suppl. Fig. 1C**). Since this amplification was indicative of the presence or absence of the cloning cassette but did not differentiate the point mutation, an alternative genotyping method which included an amplification using the EWMF (CCCTACTTTTGCCGCCTAGCTA) and EWMR Primer (CCCTACTTTTGCCGCCTAGCTA) (**Suppl. Fig. 1A**) followed by an enzymatic restriction using NheI, a restriction enzyme that can cut when the animal contains the mutation but it cannot cut in a C57Bl/6 animal (**Suppl. Fig. 1D, E**). The accuracy of this result was confirmed by sequencing the fragment included between the primers EWLPF (CTCACAGCTCTAATCCTCCTCCCTC) and EWMR (CCCTACTTTTGCCGCCTAGCTA) (**Suppl. Fig. 1A**). Finally, to also confirm the phenotype, a western blot probed against pCx43^S368^ demonstrated the absence of this phosphorylation in Cx43^S368A^ mice but not in C57Bl/6 mice (**Suppl. Fig. 1F, G**).

### Weight drop traumatic brain injury model

We used an impact acceleration weight drop model, as described previously (George et al., 2021; Shandra et al., 2019; Shandra & Robel, 2020). Briefly, mice were anesthetized using 3.5% isoflurane for five minutes (min). Immediately after and while they were still unconscious, mice were administered the analgesic buprenorphine (0.05-0.100 mg/kg) and placed on a foam pad. A metal plate was positioned over their head to create a diffuse injury and a 100 g weight was dropped from a 50 cm height. Mice were placed on a heat pad until consciousness was recovered. Sham mice (control) went through the same analgesics and anesthesia procedures but were not hit by the weight. For most experiments, three injuries with 45 min between them were inflicted. In experiments in which Cx43^S368A^ mice were used, mice were only injured twice, with 45 min between the injuries due to an increased mortality in both Sham and TBI groups.

### Western blot

Male and female mice were sacrificed by cervical dislocation at the time indicated for each experiment. Their brain was extracted and the cortices were isolated, snap frozen in liquid nitrogen, and stored at −80°C. Tissues were mechanically homogenized in 0.5 mL of lysis buffer (50 mM TrisHCl, 150 mM NaCl, 2% NP-40, 1% sodium deoxycholate acid (DOA), 0.1% SDS at pH 8.0, phosphatase inhibitors and protease inhibitors (Sigma-Aldrich, catalog #P8340, #P0044)), followed by sonication (3x 10 s pulse/rest 5 s, 30% amplitude) on ice. Samples were incubated on ice for 15 min and centrifuged for 5 min at 1400 x *g* at 4°C. The pellet was discarded, and the supernatant protein concentration was analyzed using a bicinchoninic acid assay (BCA, Pierce, ThermoFisher Cat. #23225). 15 μg of protein was mixed with sample buffer (4x Laemmli sample buffer with β-mercaptoethanol) and 1:10 dithiothreitol, denatured at 95°C for 5 min, and loaded into Bio-Rad Criterion TGX precast 18-well gels. Gels were run at 80 V until the dye line narrowed and then at 200 V for ∼30–45 min. The Trans-blot Turbo Transfer system was used for semidry transfer at 2.5 A and 25 V for 10 min. Membranes were blocked using Azure Blocking solution for 15 min and were incubated overnight at 4°C with primary antibody (**Table 1**). After 15 min in Azure washing solution at room temperature, the membranes were incubated with secondary antibodies for 45 min (**Table 1**). Secondary antibodies were conjugated to either an infrared fluorophore or to horseradish peroxidase (HRP). After incubation, the membranes were washed for 15 min at room temperature. Images were acquired using either a Li-Cor Odyssey M when infrared secondary antibodies were used or Chemidoc MP imaging system (Bio-Rad). For the latter, SuperSignal™ West Femto Maximum Sensitivity Substrate was used. Full western blots are shown **in Suppl. Figs. 3, 4, 6, 8**.

**Table 1.** Manufacturer information, working dilutions and resource identification numbers for primary and secondary antibodies that were used for western blot.

| Target protein | Manufacturer | Catalog # | RRID | Species Raised In | Monoclonal / Polyclonal | Concentration | Figure in which it was used |
| --- | --- | --- | --- | --- | --- | --- | --- |
| **Primary Antibodies** |  |  |  |  |  |  |  |
| Cx43 | Sigma-Aldrich | C6219 | AB_476857 | Rabbit | Polyclonal | 1:3000 | Fig. 1A, Fig. 2A, Fig. 4A |
| Cx43 | Millipore | MAB3067 | AB_94663 | Mouse | Monoclonal | 1:1000 | Fig. 4C |
| pCx43S368 | Thermo Fisher Scientific | 48-3000 | AB_2533845 | Rabbit | Polyclonal | 1:1000 | Fig. 4A, Fig. 4C |
| GAPDH | Abcam | ab8245 | AB_2107448 | Mouse | Monoclonal | 1:5000 | Fig. 1A, Fig. 2A, Fig. 4A |
| **Secondary Antibodies** |  |  |  |  |  |  |  |
| Rabbit IgG HRP | Santa Cruz Biotechnology | sc-2357 | AB_628497 | Mouse | Monoclonal | 1:5000 | Fig. 1A, Fig. 2A, Fig. 4A |
| m-IgGk BP-HRP | Santa Cruz Biotechnology | sc-516102 | AB_2687626 |  | Monoclonal | 1:5000 | Fig. 1A, Fig. 2A, Fig. 4A |
| Rabbit IgG IR700 | AzureSpectra | AC2128 | AB_3665331 | Goat | Unknown | 1:5000 | Fig. 4C |
| Mouse IgG IR800 | AzureSpectra | AC2135 | AB_3665332 | Goat | Unknown | 1:5000 | Fig. 4C |

Cx43 is typically observed as several bands in the vicinity of 43 kDa representing differing phospho-isoforms (P0-P3) (Musil & Goodenough, 1991). Resolution of these bands is entirely dependent on running buffer selection and electrophoresis conditions. For Cx43 western blots, 4-20% gradient gels were used to compress these Cx43 bands into a more quantifiable single band (**Suppl. Fig. 3**).

### Immunohistochemistry

Mice were anesthetized using a fatal dose of ketamine (100 mg/kg) and xylazine (10 mg/kg). When the mice were unconscious, they were transcardially perfused with PBS, followed by 4% PFA. After brain extraction, they were post-fixed in 4% PFA overnight, transferred to PBS, and coronally sliced at 50 µm thickness. Immunohistochemistry was done as described previously (George et al., 2021; Munoz-Ballester et al., 2022; Shandra et al., 2019). Briefly, 5-6 brain slices from different areas of the brain were incubated in primary antibody (**Table 2**) overnight at 4°C on a rocking platform. After washes in PBS, slices were incubated in secondary antibody (**Table 2**) with 4’,6-diamidino-2-phenylindole (DAPI, 1:1000, Invitrogen) for 45 min at room temperature on a rocking platform, washed in PBS, and mounted using Aqua-Poly/Mount (Polysciences). After drying, imaging was done using a NikonA1R confocal microscope with Apo 20x air objective and Apo 40x and 60x oil immersion objectives. The step size used during imaging was 1 µm. We imaged cortical layers I-III of the cortical gray matter excluding the piriform cortex. For each mouse, 5 images from 3-5 different slices were taken and analyzed.

**Table 2.** Manufacturer information, working dilutions and resource identification numbers for primary and secondary antibodies that were used for immunohistochemistry.

| Target protein | Manufacturer | Catalog # | RRID | Species Raised In | Monoclonal/ Polyclonal | Concentration | Figure in which it was used |
| --- | --- | --- | --- | --- | --- | --- | --- |
| **Primary Antibodies** |  |  |  |  |  |  |  |
| Cx43 | Sigma-Aldrich | C6219 | AB_476857 | Rabbit | Polyclonal | 1:1000 | Fig. 1C, D; Fig. 2E; Fig. 5A; Suppl Fig. 1 |
| Cx43 | Abcam | ab87645 | AB_1951148 | Goat | Polyclonal | 1:250 | Suppl Fig. 1 |
| S100β | Sigma-Aldrich | S2532 | AB_477499 | Mouse | Monoclonal | 1:1000 | Fig. 3 |
| **Secondary Antibodies** |  |  |  |  |  |  |  |
| Rabbit Alexa-488 | Jackson Immuno Research | 111-546-144 | AB_2338057 | Goat | Polyclonal | 1:1000 | Fig. 1C, D; Fig. 2E; Fig. 5A; Suppl Fig. 1 |
| Mouse Alexa-647 | Jackson Immuno Research | 115-606-003 | AB_2338921 | Goat | Polyclonal | 1:1000 | Fig. 3 |
| Goat Alexa-488 | Jackson Immuno Research | 705-545-003 | AB_2340428 | Goat | Polyclonal | 1:1000 | Suppl Fig. 1 |

### Triton X-100 Solubility assay

Snap-frozen tissue samples were weighed and added to 1% Triton X-100 buffer (50 mM Tris pH 7.4, 1% Triton X-100, 2 mM EDTA, 2 mM EGTA, 250 mM NaCl, 1 mM NaF, 0.1 mM Na_3_VO_4_; supplemented with Halt protease and phosphatase inhibitor cocktail, Thermo Fisher) at a final concentration of 100 mg/ml. Samples were homogenized using a bead mill and nutated for 1 h at 4°C. At this point, 10% of the lysate was removed and 4X LDS NuPAGE sample buffer (Thermo Fisher) supplemented with 400 mM DTT was added before sonication and centrifugation at 10,000 x *g* for 20 min to obtain the ‘total’ fraction. The remaining lysate was fractionated by centrifugation for 30 min at 15,000 x *g* in pre-weighed microcentrifuge tubes. The supernatant was removed and combined with 4x LDS NuPAGE sample buffer before sonication and centrifugation at 10,000 x *g* for 20 min to obtain the ‘soluble’ fraction. Pellets were weighed and suspended in 1x LDS NuPAGE sample buffer to a final concentration of 30 mg/ml. Pellets were solubilized with disruption by serial passage through an insulin syringe and sonication before centrifugation at 10,000 x *g* for 20 min. Protein samples were denatured by heating at 70 °C for 10 min and underwent SDS-PAGE with NuPAGE Bis-Tris 4-12% gradient gels and MES running buffer (Thermo Fisher). Proteins were transferred to LF-PVDF membranes (Bio-Rad), with membranes methanol-fixed and air-dried post-transfer. Following methanol reactivation, membranes were blocked in 5 % nonfat milk (Carnation) in TNT buffer (0.1 % Tween 20, 150 mM NaCl, 50 mM Tris pH 8.0) for 1 h at room temperature. Primary antibody incubation was conducted overnight at 4 °C using rabbit anti-Cx43 (1:4,000; Sigma-Aldrich) and mouse anti-GAPDH (1:500; Santa Cruz Biotechnology). Membranes were rinsed twice and washed 3 x 10 min in TNT buffer. Secondary antibody labeling was conducted for 1 h at room temperature using antibodies conjugated to Alexa Fluor 555 and Alexa Fluor 647 (Thermo Fisher). Following the secondary antibody incubation, membranes were washed in TNT buffer as previously described. Prior to imaging, membranes were fixed in methanol and air-dried. Images were acquired using a Chemidoc MP imaging system (Bio-Rad). Western blot quantification was conducted using Bio Rad Image Lab software.

### Super-resolution imaging

Super-resolution imaging was performed in cortical layers II/III of the somatosensory cortex in 50 µm coronal brain slices using a Nikon AX R Confocal Microscope System with Nikon Spatial Array Confocal (NSPARC). Images were acquired with a 60x objective and 5x digital zoom. Imaging was conducted using a Galvano scanner to capture z-stacks. The initial plane of focus was identified at the top of each tissue slice from which 1 µm thick z-stacks were taken with a step size of 0.1 µm. 15 z-stacks were captured per mouse in 3 different brain sections (5 z-stacks per section). Imaging parameters were applied consistently throughout the experiment. For data in Fig. 3, Z-stacks were deconvolved in Nikon NIS-Elements software using blind deconvolution with 10 iterations. For analysis, the fourth plane from the top of each slice was selected.

### Ethidium bromide assay

Three days after inducing TBI, mice were sacrificed by cervical dislocation and their brains were rapidly removed and immersed in ice-cold cutting solution (in mM): 135 N-methyl-D-gluoconate (NMDG), 1.5 KCl, 1.5 KH_2_PO_4_, 23 choline bicarbonate, 25 D-glucose, 0.5 CaCl_2_ 2.5 MgSO_4_. Slices of 200 µm thickness were cut using a vibratome and placed in a 6-well plate with hanging cell culture inserts, one for each condition, in carbogen-bubbled artificial cerebrospinal fluid (ACSF) (in mM): 125 NaCl, 3 KCl, 1.25 NaH_2_PO_4_, 25 NaHCO_3_, 2 CaCl_2_, 1.3 MgSO_4_, 25 D-glucose at room temperature. After 10 min recovery in ACSF, a different treatment was added to each well and incubated for 15 min. The treatments were: water, DMSO, 200 µM carbenoxolone disodium salt (CBX) (Sigma-Aldrich), 10 µM spironolactone (Sigma-Aldrich), and 2 µM the hemichannel inhibitor peptide TAT-Gap19 (YGRKKRRQRRRKQIEIKKFK, synthetized by Genscript). Slices in water were used as a control for CBX and TAT Gap19 slices. DMSO was used as a control for the spironolactone treatment. Following the treatment incubation, 4 µM ethidium bromide (EtBr) was added to the well and incubated for 15 min, followed by a 10-minute wash in ACSF. Slices were fixed in 4% PFA for 20 min. After that, immunohistochemistry against S100β was performed as described in the immunohistochemistry section, followed by confocal imaging of S100β/EtBr+ astrocytes.

### Seizure susceptibility assay using sub-convulsive dosing of Pentylenetetrazole

To determine seizure susceptibility, C57Bl/6 and Cx43^S368A^ TBI, Sham (all procedures as in TBI mice including administration of isoflurane and buprenorphine except for the weight drop) or naïve mice (no isoflurane or buprenorphine) were injected with sub-convulsive doses of Pentylenetetrazole (PTZ) on alternating days, starting on day 3 following TBI/Sham. Naïve C57Bl/6 and Cx43^S368A^ mice were injected 8–11 times and TBI/Sham C57Bl/6 and Cx43^S368A^ mice were injected 11 times. PTZ was made freshly for each session at 3.5 mg/mL in saline followed by intraperitoneal administration of 10 µL/g body weight (35 µg/g body weight). Following PTZ injection, mice were observed for 30 min and then returned to their home cage. Behavioral abnormalities indicating seizures were scored according to a modified Racine scale (Dhir, 2012). In short, a score of 0 indicated no behavioral abnormalities, freezing behavior and/or laying down were scored as 1, shaking and/or twitching were scored as 2, forelimb clonus, lordotic posture and tail erection were scored as 3, rearing and falling were scored as 4 and generalized, tonic-clonic seizures, jumping, and loss of postural tone sometimes resulting in death were scored as 5. The time to onset of generalized, tonic-clonic seizures was recorded. One mouse died 5 seconds after the injection and was excluded from the study given that mortality was likely due to an inappropriate injection site.

### Data analysis

#### Western Blot quantification

Western blot images were imported as a third-party image and analyzed using Image Studio software (Li-Cor) (**Fig. 1, 4**) or ImageLab (BioRad) (**Fig. 2**). Cx43 or pCx43^S368^ bands were identified using the automatic *Add rectangle* function of Studio Image, and rectangular volume analysis in ImageLab. For the solubility assay, insoluble (junctional) fraction values were expressed relative to soluble (non-junctional) values from the same samples. The background was calculated based on the median intensity at the top and bottom area surrounding the band (3 pixels). The normalized value of the signal based on the background provided by the program was used as the intensity value. GAPDH or total protein was used as an internal loading control, as specified for each experiment.

**Figure 1.**
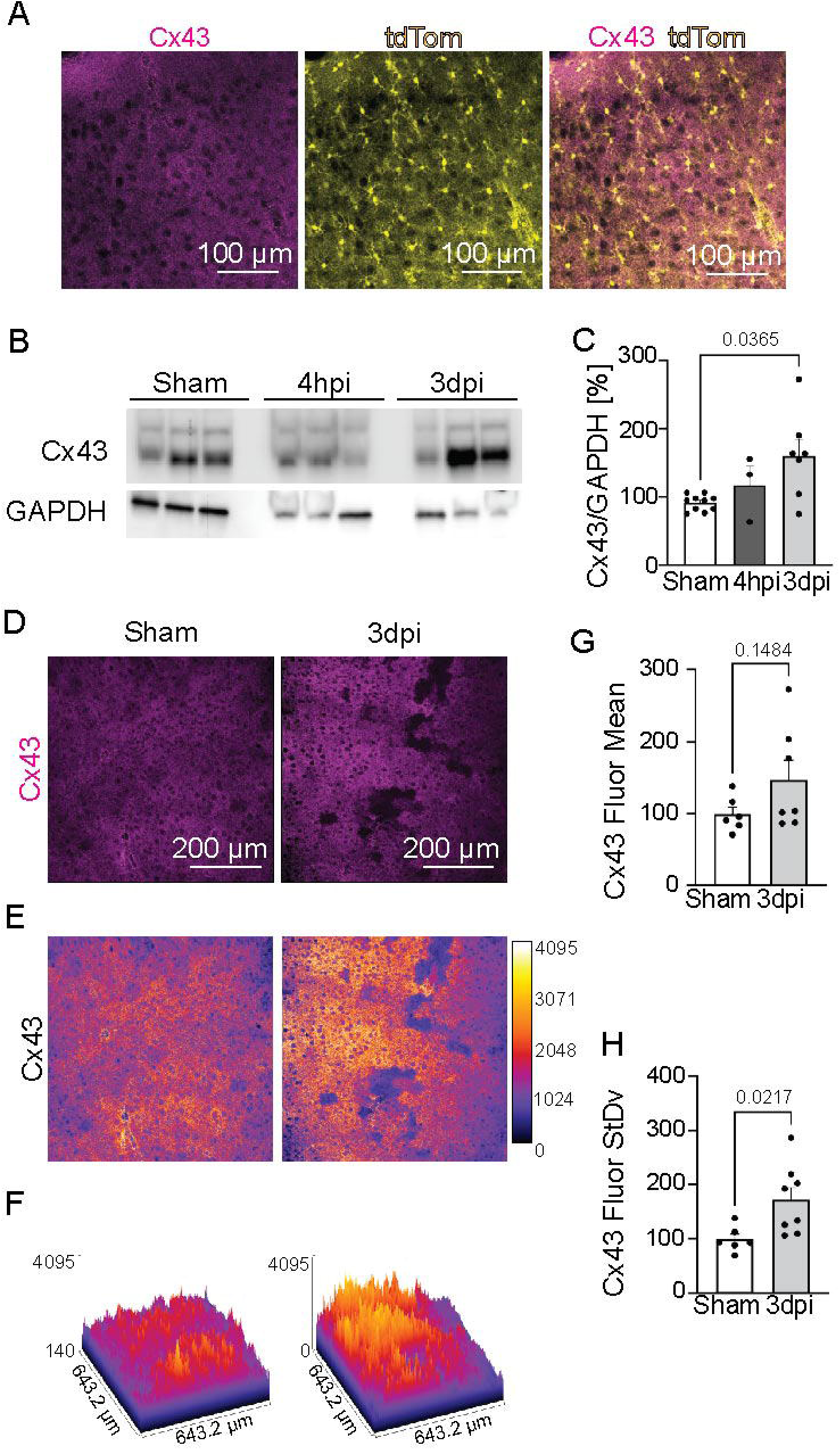
Mild TBI induces a heterogenous response in Cx43 protein expression. **A** Representative immunohistochemistry image of cortical tissue demonstrating Cx43 colocalization with Aldh1l1-tdTomato-positive astrocytes. **B** Representative western blot of cortical brain samples probed against Cx43 protein in Sham, 4 hpi and 3 dpi mice. Each lane represents a different animal. GAPDH was used as loading control. Representative western blots presented here were cropped from a membrane with additional time points (**Suppl. Fig. 3**). **C** Quantification of Cx43 protein levels normalized to GAPDH. **D** Representative Cx43 immunohistochemistry images of cortical tissue. **E** Quantification of mean fluorescence intensity of immunohistochemistry images. **F** Same image as in **1D** in which each pixel is colored based on its intensity value. **G** Surface plot of the images in figure 1D in which each pixel color and height indicates the intensity level of said pixel. **H** Normalized means of the standard deviation values for the intensity fluorescence of Cx43 immunohistochemistry images. In all figures, group statistics were run after averaging data per mouse. Data points in scatterplots represent individual mice and means were displayed as bar graphs with standard error of the mean.

**Figure 2.**
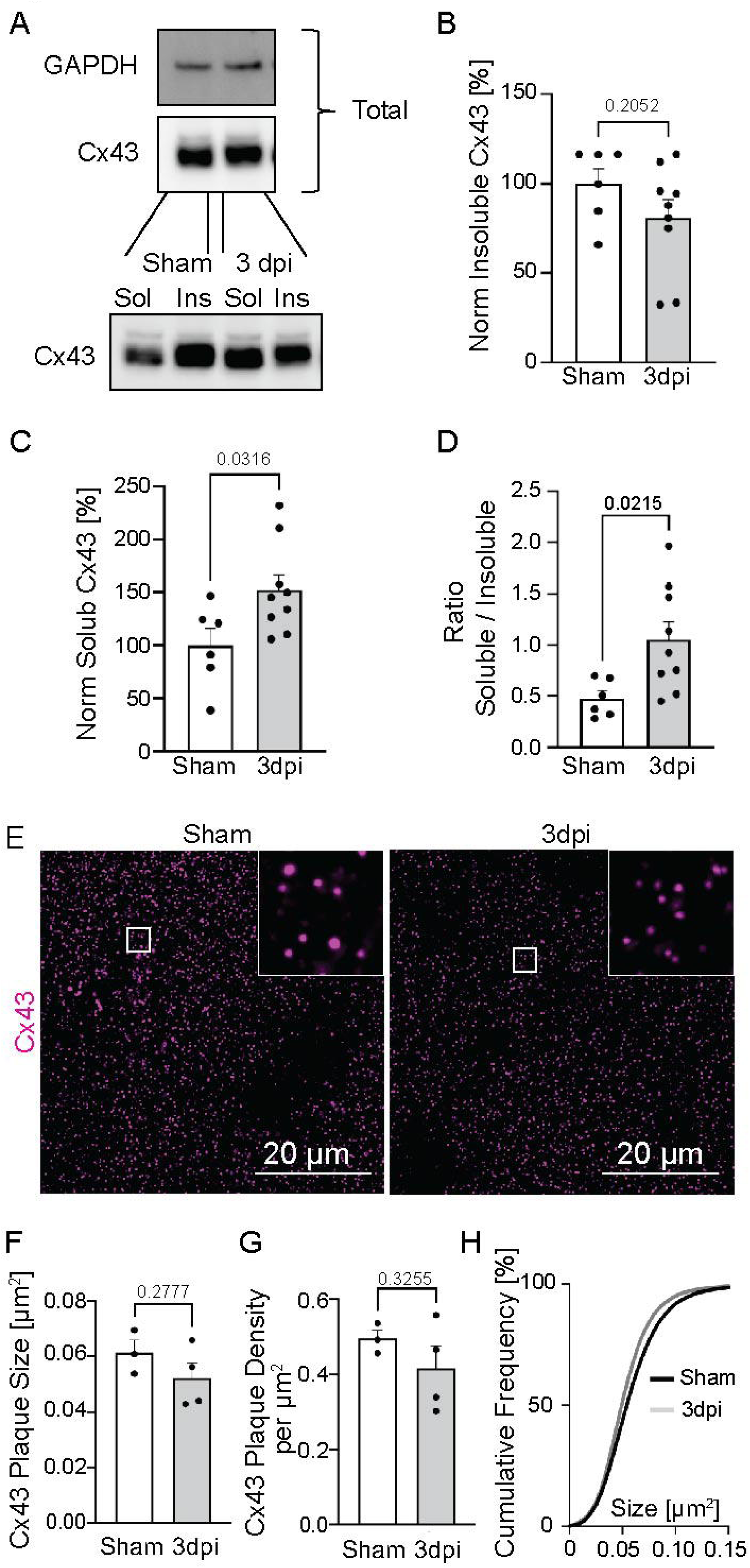
Cx43 gap junction plaque density and size are unaffected by mild TBI while soluble Cx43 increases. **A** Representative western blot of solubility-fractionated cortical gray matter tissue. Sol: soluble fraction, Insol: insoluble fraction. **B** Quantification of the Cx43 insoluble fraction. **C** Quantification of the Cx43 soluble fraction. **D** Ratio of soluble to insoluble Cx43 fractions. **E** Representative super-resolution images of Cx43 plaques. White square indicates area zoomed and their location in the global image. **F** Quantification of Cx43 plaque size in Sham and 3 dpi. **G** Quantification of Cx43 plaque density in Sham and 3 dpi. **H** Graph of the cumulative distribution of Cx43 plaque size, black line = Sham and grey line = 3 dpi.

#### Cx43 fluorescence intensity quantification

The mean, standard deviation, and kurtosis of the Cx43 signal were calculated per image using the *Measurement* function of Fiji (ImageJ). The value of each slice was averaged per mouse. The average of Sham mice was considered 100%, and each value was normalized based on that. Images containing the meninges were excluded from the analysis since meningeal fibroblasts express high levels of Cx43 and bias results.

#### Cx43 super-resolution measurements of plaque size and density

General Analysis 3 imaging processing and measuring module in the Nikon imaging software, NIS-Elements, was used to threshold super-resolution images to eliminate background signal before binarization to isolate the Cx43 signal. Within NIS-Element’s General Analysis 3, ObjectCount, ObjectArea, and MeasuredArea functions were used to measure plaque number, plaque size, and image area, respectively. Average plaque size was determined by the sum of all plaque areas divided by the total number of plaques per mouse. Average density was determined by the total number of plaques divided by the total area imaged per mouse.

#### EtBr fluorescence intensity analysis

For each image, astrocytes were blindly identified using S100β staining and delimited using the *Oval selection* feature of ImageJ. After that, the EtBr mean fluorescence intensity within that circle was calculated using the *Measurement* function of FIJI. The mean fluorescence intensity of an oval selection of the tissue with no signal was used as background and subtracted from the mean fluorescence intensity values of that slice. Each value was averaged per slice, and each slice was averaged per mouse. Per animal, 3-5 slices were analyzed after taking 5 images per slice. In each image 5 randomly selected astrocytes were analyzed. For the comparison between Sham and 3 dpi mice, the average fluorescence intensity of Sham mice was considered 100%, and each mouse value was normalized accordingly. For the comparison between control and treatment with inhibitors within 3 dpi mice, the average fluorescence intensity of the control treatment was considered 100%, and all the values were normalized based on this value. In one batch of experiments, CBX was expired, no inhibition was observed in any sample, and those values were excluded from the analysis.

### Statistical analysis

Statistics were performed and graphed using GraphPad Prism 9 or 10 (GraphPad Software). Data were tested for normality using the Kolmogorov–Smirnov (KS) normality test. Before running ANOVA tests, homoscedasticity was tested using the Levene test. If the data did not meet the conditions of applicability for parametric tests, non-parametric tests were used. Statistical tests were chosen accordingly and are specified for each experiment in the results section. In each dataset, the outlier test ROUT at Q 1% was run, and values identified as outliers were excluded from the analysis. Two-tailed statistical tests were used unless the prediction was unidirectional. In this case, one-tailed tests were used. Post-hoc analysis was used when there were more than two groups. The specific test that was used is stated for each experiment either in the results section or figure legend. All group statistics were run after averaging data per mouse. Data points in scatterplots represent individual mice and means were displayed as bar graphs with standard error of the mean. Exact p-values are reported within the results section and figures. While both sexes were used in the study, the study was not powered to assess sex differences, and data from mice of both sexes were combined.

## Results

### Mild TBI induces heterogeneous changes in astrocytic Cx43

In the adult cortical gray matter, Cx43 is most abundantly expressed in astrocytes and–to a much lesser extent–in some vascular cell types (Magnotti et al., 2011; Zhan et al., 2023). Thus, as expected, immunohistochemistry against Cx43 in uninjured Sham mice demonstrated localization of the majority of Cx43 protein in fine astrocytic processes and in astrocyte endfeet along blood vessels (**Fig. 1 A, Suppl. Fig. 2**) and very little overlap with CD31-positive endothelial cells (**Suppl. Fig. 2**).

To determine temporal and spatial changes in Cx43 protein levels after mild TBI, mice were subjected to weight drop injury, which induces diffuse brain injury across both hemispheres, as previously described (Abd-Elfattah Foda & Marmarou, 1994; Marmarou et al., 1994; Nichols et al., 2016). When calibrated to mild TBI, this diffuse injury occurs in the absence of focal lesions (no lesioned area or injury focus can be defined within the tissue), primary cell death, or astrocyte border/scar-formation within the cortical gray matter (Munoz-Ballester et al., 2022; Shandra et al., 2019). Cortical Cx43 protein levels detected by western blot (WB) did not change 4 hpi (4hpi mean = 117.5±27.70) when compared to Sham mice (Sham mean = 91.86±3.622; Kruskal-Wallis multiple comparisons test Sham vs 4hpi p-value = 0.9966). However, a significant increase in Cx43 protein was observed 3 dpi (3 dpi mean = 160.7±23.82; Kruskal-Wallis multiple comparisons test Sham vs 3 dpi p-value=0.0365) (**Fig. 1 B,C, Suppl. Fig. 3**).

When immunohistochemistry against Cx43 was performed after mild TBI/Sham, the Cx43 expression pattern was heterogeneous at 3 dpi: while some areas had a decrease in Cx43, other areas appeared to have increased levels of Cx43 (**Fig. 1 D-F).** Surface plots in which the height of a pixel is proportional to the intensity of this pixel reflected a high heterogeneity in the Cx43 fluorescence intensity across different astrocyte populations at 3 dpi when compared to Sham (**Fig. 1 F**). While Cx43 mean fluorescent intensity was not significantly changed (Sham mean = 99.30±9.866, n = 6; 3 dpi mean = 147.4±27.24, n = 7; unpaired two-tailed t-test p-value = 0.1484) (**Fig. 1 E**), the variability of the samples measured as standard deviation was higher at 3 dpi when compared to Sham (Sham mean = 99.17±9.687, n = 6; 3 dpi mean = 69.83±22.59, n = 8; unpaired two-tailed t-test, p-value = 0.0217) (**Fig. 1 H**) pointing to a regional heterogenous response. We conclude that mild TBI induces a heterogenous astrocyte response, in which Cx43 is locally regulated, being downregulated in a subset of astrocytes and increased in a different subset. This is in alignment with previous studies of astrocytes in this model in which astrocytes respond heterogeneously to mild TBI across the cortical gray matter (George et al., 2021; Shandra et al., 2019).

### Mild TBI leaves mean Cx43 gap junction density unaffected while soluble Cx43 increases

To determine if the overall increase in Cx43 is reflected in an increase in Cx43 GJs, we performed a fractionation assay based on Cx43 Triton X-100 solubility and analyzed the fractions using western blot. Species of Cx43, which remain in the ‘soluble’ fraction, encompass monomeric Cx43 from the cell interior, connexons enroute to and from the plasma membrane, and hemichannels on the cell surface. GJs are not solubilized by the non-ionic Triton X-100 detergent due to the double membranous nature of these structures, and they are fractionated out by centrifugation to the pellet, enabling quantification of junctional (insoluble) vs. non-junctional (soluble) Cx43 (Musil & Goodenough, 1991) (**Fig. 2 A; Suppl. Fig. 4)**. While no change was observed in the amount of Cx43 within the Triton X-100 insoluble fraction (GJs) (Sham mean = 100±8.601, n = 6; 3 dpi mean = 80.91±10.11, n=9; unpaired two-tail t-test p-value = 0.2052) (**Fig. 2 B**), Cx43 levels in the Triton X-100 soluble fraction (non-junctional Cx43) increased at 3 dpi (Sham mean = 100±15.81, n=6; 3 dpi mean = 152.5±14.30, n=9; unpaired two-tail t-test p-value = 0.0316) (**Fig. 2 C**). Accordingly, the ratio of soluble to insoluble Cx43 was increased (Sham mean = 0.4744±0.07403, n=6; 3 dpi mean = 1.056±0.1729, n=9; unpaired two-tail t-test p-value = 0.0215) (**Fig. 2 D**).

We next used super-resolution imaging microscopy to determine whether Cx43 plaques changed in density or size. Plaques are routinely defined by confocal microscopy as dense accruals of Cx43 immunostaining at cell-cell borders and encompass functional or closed GJs. Hemichannels are less amenable to detection by confocal microscopy given their diffuse distribution across the plasma membrane (Lucaciu et al., 2024). After processing the images to isolate junctional plaques, no difference in plaque size was found between Sham and 3 dpi mice (Sham mean = 0.06139 µm^2^ ±0.00456, n=3; 3 dpi mean = 0.05226 µm^2^ ±0.005454; unpaired two-tail t-test p-value = 0.2777) or plaque density was found between Sham and 3 dpi mice (Sham mean = 0.4951 plaque/µm^2^ ±0.02272, n = 3; 3 dpi mean = 0.4167 plaque/µm^2^ ±0.05862, n=4; unpaired two-tail t-test, p-value = 0.3255) **(Fig. 2 E–H**), suggesting that GJ turnover is not affected by TBI at 3 dpi. These results indicate that the Cx43 increase after mild TBI does not lead to increased GJ formation but instead leads to an increase in soluble, non-junctional Cx43 composed of hemichannels and/or cytoplasmic Cx43.

### Mild TBI may cause an increase in hemichannel activity

To assess whether the increase in Cx43 expression was due to an increase in Cx43-dependent hemichannels, we used an ethidium bromide (EtBr) uptake assay in acute slices from Sham and TBI mice at 3 dpi. EtBr, a DNA intercalating agent, can pass through open Cx43 hemichannels or pannexin channels on the cell surface and subsequently be detected in the UV or far-red channel. Cx43 hemichannel activity of a cell can be inferred based on the fluorescence intensity of its nucleus because this fluorescence is proportional to the activity of the channel through which it enters (Contreras et al., 2003; Hansen et al., 2014; Johnson et al., 2016). We labeled astrocytes using an antibody against S100β in Sham and 3 dpi mice. Nuclei of S100β-positive astrocytes had increased EtBr fluorescence intensity at 3 dpi compared to Sham mice (**Fig. 3 A,B**) (Sham Mean = 100.0±2.036, n= 4; 3 dpi Mean = 185.8±14.91, n=6, one-tailed unpaired t-test p-value = 0.0009).

**Figure 3.**
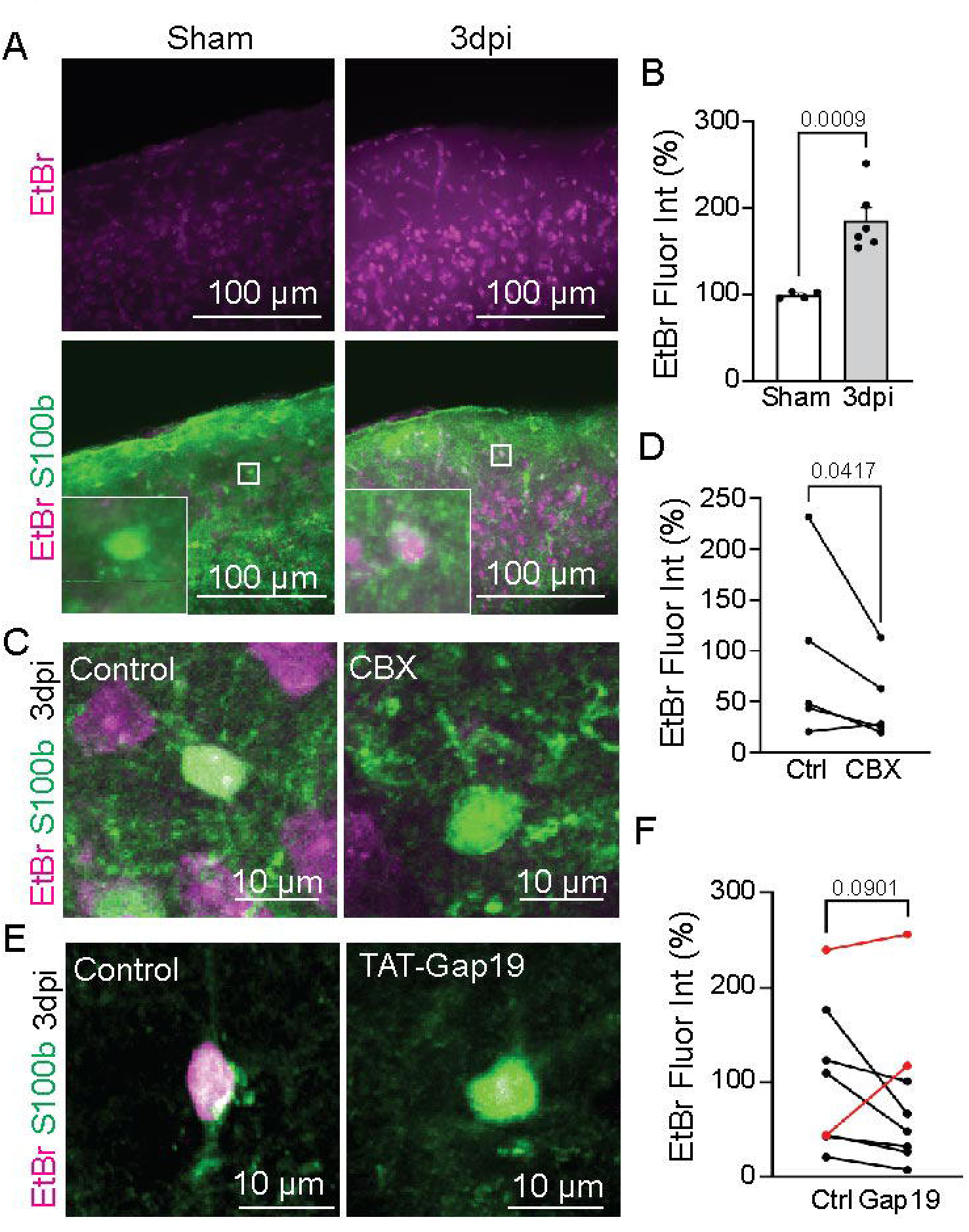
Mild TBI induces an increase in Cx43 hemichannel-dependent activity. **A** Representative confocal image of Sham and 3 dpi acute brain slices after EtBr (magenta) uptake assay and immunohistochemistry for S100β (astrocyte marker) in green. Upper panels: EtBr signal alone. Lower panels: merge imaged of EtBr and S100β. White squares indicate the zoomed area. **B** Quantification of EtBr fluorescence intensity levels in acute brain slices of Sham and 3 dpi mice. **C** Representative confocal images of control and CBX-treated acute slices at 3 dpi. **D** Quantification of EtBr fluorescence intensity levels after CBX treatment. **E** Representative confocal images of a control and a TAT-Gap19-treated acute slices at 3 dpi. **F** Quantification of EtBr fluorescence intensity levels after TAT-Gap19 treatment. Red dots and lines indicate the samples that show no decrease in EtBr fluorescence intensity.

To determine if the increase of the EtBr signal was specific to hemichannels, we first used carbenoxolone (CBX), a broader GJ/hemichannel/pannexin blocker (Sagar & Larson, 2006). As predicted, EtBr uptake was significantly reduced in acute brain slices 3 dpi mice when compared to Sham (Sham mean = 90.97 ± 38.22, n=5; mean CBX = 50.00 ± 17.45; paired one-tail test p-value = 0.0417) **(Fig. 3 C,D**). We next assessed more specific inhibitors to determine if EtBr uptake is dependent on pannexins or Cx43 hemichannels. We next used the Cx43 hemichannel-specific inhibitor TAT-Gap19. EtBr uptake detected in TBI mice was on average decreased by 20% when TAT-Gap19 was used, although this difference did not reach the significance threshold (Sham mean = 100 ± 27.34, n=8; TAT Gap19 mean = 81.74 ± 28.18, n=8; paired one-tail t-test analysis, p=0.0901). However, in pairwise comparison of slices from the same mouse 6 of 8 mice treated with vehicle or TAT-GAP19 showed a reduction in EtBr uptake after blocking Cx43 hemichannel function (**Fig. 3 E,F**). In contrast, the pannexin blocker spironolactone (10 µM) did not reduce astrocytic EtBr (Control mean = 100.0 ± 33.80, n=6; spironolactone mean = 123.7 ± 24.93, n=6; paired one-tail test p-value = 0.1678). Pairwise assessment of slices from the same mouse treated with either vehicle or spironolactone suggests an increase of EtBr intensity in 4 of 6 mice in response to the pannexin inhibitor (**Suppl Fig. 5**). Thus, pannexin channels are not responsible for EtBr uptake into astrocytes after mild TBI. In conclusion, EtBr uptake is increased in cortical astrocytes after mild TBI. Notably, while dying cells may take up EtBr, we previously demonstrated an absence of cell death in this model of mild TBI (Shandra et al., 2019). Further, unspecific EtBr uptake would be unaffected by CBX, which effectively blocked EtBr uptake into astrocytes after mild TBI. Thus, Cx43 hemichannel contribution to this uptake occurred in 75% of the mice while the other 25% responded differently, possibly indicating molecular mechanisms underlying the well-known heterogeneity in TBI outcomes from patient to patient, if not due to variable efficacy or diffusion of TAT-Gap19 in thick acute slices.

### Cx43 phosphorylation at serine 368 impacts seizure susceptibility

Phosphorylation at Cx43^S368^ (pCx43^S368^) is well described in regulation of Cx43 dynamics and GJ formation, function, and stability (Calhoun et al., 2020; Ek-Vitorin et al., 2006; Lampe et al., 2000; Sirnes et al., 2009). To determine if this post-translational modification was associated with the Cx43 alterations we described above after mild TBI, we assessed the levels of pCx43^S368^ by western blot and found an increase at 3 dpi (Sham mean = 100.0±23.48, n=6; 3 dpi mean = 189.0 ±23.01, n=5; two-tailed t-test p-value = 0.0253) (**Fig. 4 A,B, Suppl. Fig. 6**).

**Figure 4.**
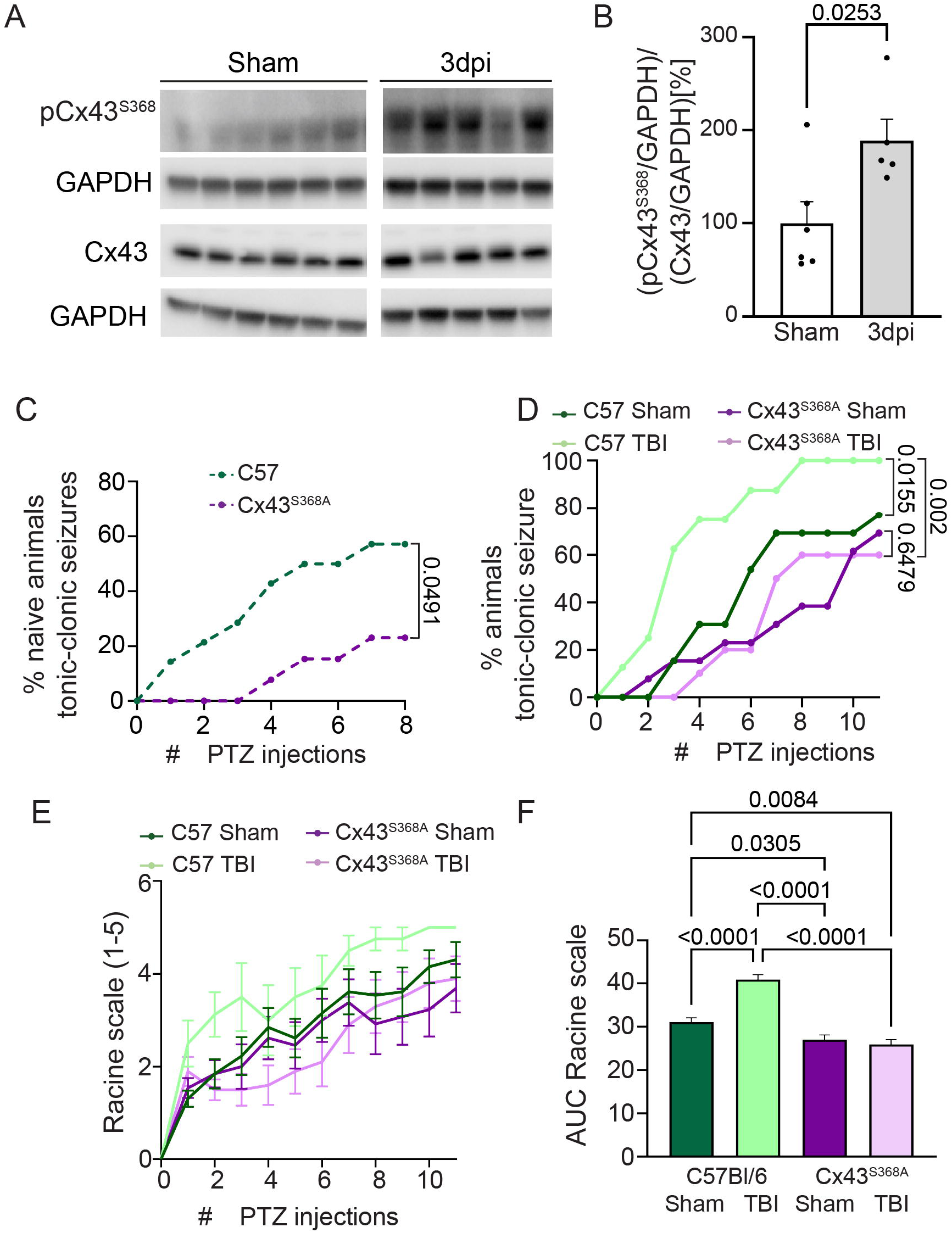
Phosphorylation of Cx43^S368^ after mild TBI increases seizure susceptibility. **A** Representative western blot of cortical brain samples of Sham and 3 dpi mice against pCx43^S368^ and Cx43 proteins. Two gels with the same samples were run to probe against pCx43^S368^ and Cx43, respectively. As an internal loading control, membranes were probed against GAPDH. Each lane represents a different mouse. **B** Quantification of pCx43^S368^ as a ratio between pCx43^S368^ normalized to GAPDH and Cx43 normalized to GAPDH. **C** The graph shows the cumulative percentage of naïve C57Bl/6 and Cx43^S368A^ mice that experienced a tonic-clonic seizure over time. Kaplan-Meier survival analysis was performed with the statistical comparison between groups indicated by p-values in the graph. C57Bl/6 n = 14, Cx43^S368A^ n = 13. **D** Graph representing the cumulative percentage of Sham C57Bl/6 (n=13), 3 dpi C57Bl/6 (n=8), Cx43^S368A^ Sham (n=13), Cx43^S368A^ and 3 dpi Cx43^S368A^ (n=10) mice which experienced a tonic-clonic seizure Kaplan-Meier survival analysis was performed with the statistical comparison between groups indicated by p-values in the graph. **E** Graph representing the average Racine scale value of each experimental group at every injection number. Sham C57Bl/6, C57Bl/6, Cx43^S368A^ Sham, Cx43^S368A^ 3 dpi. **F** Quantification of the area under the curve of the graph in E.

We next used Cx43^S368A^ mice harboring a serine-to-alanine substitution within the Cx43^S368^ site (**Suppl. Fig. 1 A**), which was designed to prevent phosphorylation of Cx43 at this site. We validated the animal model by sequencing the Cx43^S368A^ allele and developed polymerase chain reaction (**Suppl. Fig. 1 B,C**) and restriction enzyme (**Suppl. Fig. 1 D,E**) genotyping protocols to confirm the serine-to-alanine substitution. We next confirmed by western blot that pCx43^S368^ was almost completely absent (C57Bl/6 mean = 100.0±6.766, Cx43^S368A^ mean = 5.881±1.341) (**Suppl. Fig. 1 F,G**).

We then asked whether pCx43^S368^ impacts seizure susceptibility in the context of TBI. We assessed seizure threshold by administering subconvulsive doses of the GABA A receptor antagonist pentylenetetrazol (PTZ) on alternate days for 2-3 weeks to C57Bl/6 and Cx43^S368A^ mice. At this dosage, PTZ has a cumulative effect that over time increases the likelihood that a mouse seizes after injection. First, we compared uninjured, naïve C57Bl/6 and Cx43^S368A^ mice (also on C57Bl/6 background). We assessed behavioral abnormalities and seizure severity using Racine scale scores 1 through 5. Surprisingly, some C57Bl/6 mice already had Racine scale 5 tonic-clonic seizures after the first injection, while Cx43^S368A^ mice had Racine scale 5 tonic-clonic seizures at the earliest after 4 PTZ injections. After 6 injections, 50% of the C57Bl/6 mice had Racine scale 5 seizures while only 15% of the Cx43^S368A^ mice did (**Fig. 4C, Suppl Table 1**).

Next, we assessed seizure susceptibility in the context of mild TBI. As expected, TBI lowered the seizure threshold (**Fig. 4D**) resulting in 62% of C57Bl/6 mice with mild TBI having Racine scale 5 seizures after the third injection, while only 15% of C57Bl/6 Sham mice did at this point. It took 8 injections for 54% of C57Bl/6 Sham mice to present with Racine scale 5 seizures. C57Bl/6 TBI mice also had higher overall Racine scores than Sham mice across all time points (**Fig. 4 E,F**). In contrast, Cx43^S368A^ mice were remarkably protected from the development of seizures compared to C57BL/6 after TBI when assessing the percentage of mice with Racine scale 5 seizures (**Fig. 4D**) or Racine scores (**Fig. 4 E,F**). It took 8 injections for 60% of Cx43^S368A^ TBI mice to display Racine scale 5 seizures. After 10 injections, all C57Bl/6 mice had tonic-clonic seizures, while only 60% of Cx43^S368A^ did by the end of the experiment after 11 PTZ injections (Area Under the Curve: C57Bl/6 Sham mean = 31.08±0.9735, n=13; C57Bl/6 3dpi mean = 40.88±1.147, n= 8; Cx43^S368A^ Sham mean = 27.00±1.109, n= 13; Cx43^S368A^ Sham 3dpi = 25.95±1.084, n= 10; two-tailed one-way ANOVA test p-value ANOVA = <0.0001; p-values for Tukey’s post-hoc multiple comparisons are indicated in **Fig. 4F**). In summary, preventing pCx43^S368^ conferred protection from the development of PTZ-induced seizures in the context of mild TBI.

### Gap junction plaque density is reduced in Cx43^S368A^ mice

To determine the mechanism by which a lack of pCx43^S368^ confers protection from seizure susceptibility, we used super-resolution imaging microscopy to assess Cx43 plaque size and density in Cx43^S368A^ mice compared to C57Bl/6 controls (**Fig. 5A**) as assessed with panCx43 immunohistochemistry. These findings are consistent when using the Cx43 staining protocol performed in **Fig. 2 E-H**, which used only one Cx43 antibody instead of a cocktail of three antibodies (**Suppl Fig. 7**).

**Figure 5.**
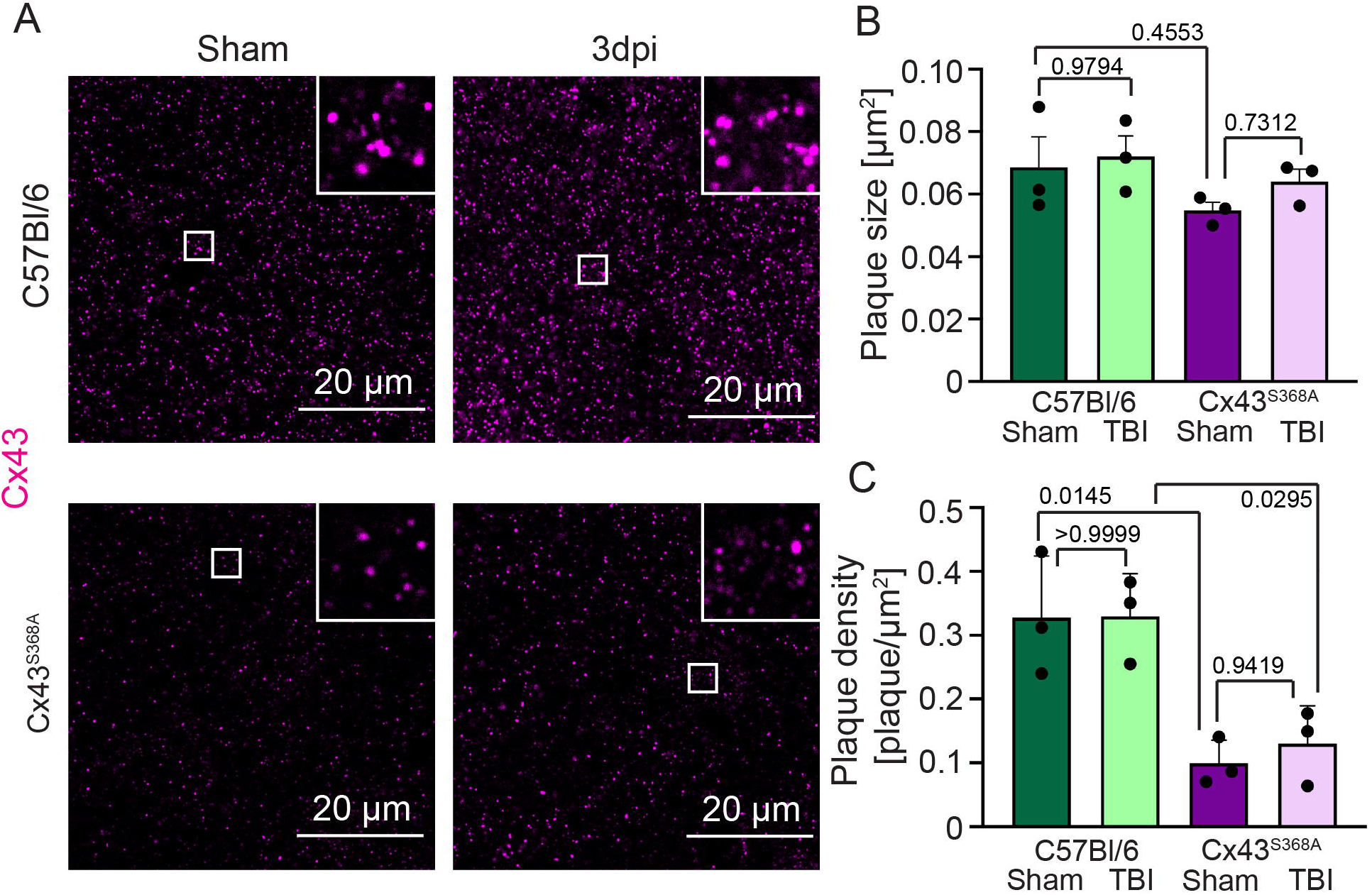
Gap junction plaque density is reduced in Cx43^S368A^ mice. **A** Representative immunohistochemistry of super-resolution images of Cx43 plaques in C57Bl/6 Sham, C57Bl/6 3 dpi, Cx43^S368A^ Sham, and Cx43^S368A^ 3 dpi mice. **B** Quantification of Cx43 plaque size in C57Bl/6 Sham, C57Bl/6 3 dpi, Cx43^S368A^ Sham and Cx43^S368A^ 3 dpi mice. **C** Quantification of Cx43 plaque density in C57Bl/6 Sham, C57Bl/6 3 dpi, Cx43^S368A^ Sham and Cx43^S368A^ 3 dpi mice.

We observed no difference in plaque size between uninjured Cx43^S368A^ and C57Bl/6 mice (C57Bl/6 Sham mean = 0.06860 µm^2^ ±0.097370, n = 3; Cx43^S368A^ Sham mean = 0.05475 µm^2^ ±0.002589, n = 3; Tukey’s multiple comparisons test p-value = 0.4553) (**Fig. 5B**), but a significant reduction in plaque density between uninjured Cx43^S368A^ and C57Bl/6 mice (C57Bl/6 Sham mean = 0.3277 plaque/µm^2^ ±0.05569, n = 3; Cx43^S368A^ Sham mean = 0.09919 plaque/µm^2^ ±0.02115, n = 3; mean Tukey’s multiple comparisons test p-value = 0.0145) (**Fig. 5C**).

When comparing Cx43^S368A^ mice with and without mild TBI, we found no difference in plaque size (Cx43^S368A^ Sham mean = 0.05475 µm^2^ ±0.002589, n=3; Cx43^S368A^ 3 dpi mean = 0.06407 µm^2^ ±0.003897, n = 4; Tukey’s multiple comparisons test p-value = 0.7312) (**Fig. 5B**) or plaque density (Cx43^S368A^ Sham mean = 0.09919 plaque/µm^2^ ±0.02115, n = 3; Cx43^S368A^ 3 dpi mean = 0.1302 plaque/µm^2^ ±0.03416, n=3; Tukey’s multiple comparisons test p-value = 0.9419) (**Fig. 5C**).

Lastly, because astrocytes also express the gap junction protein Cx30, which may compensate for changed Cx43 (Theis et al., 2003), we assessed Cx30 protein levels and plaques to determine if the modification to Cx43 affects Cx30 levels. We found no significant differences in Cx30 protein levels or plaque densities between Cx43^S368A^ and C57Bl/6 mice (**Suppl. Fig. 8**) suggesting that Cx30 does not compensate.

These results demonstrate reduced gap junction plaque densities in mice that are unable to phosphorylate Cx43^S368^. Although future studies need to address whether astrocytes are more or less coupled in Cx43^S368A^ mice (i.e. if more or less GJs are open), this data suggests that a lack of pCx43^S368^ confers protection from the development of hyperexcitability and seizures and may be a potential therapeutic target.

## Discussion

Despite substantial literature reporting altered Cx43 regulation in brain pathology, there is still limited knowledge regarding the role of Cx43 and/or gap junction regulation in TBI, especially mild TBI/concussion. In many studies, the expression of Cx43 mRNA or protein is interpreted as equivalent to gap junctional coupling (Deshpande et al., 2017; Garbelli et al., 2011) without consideration of the non-junctional roles of Cx43. Yet, Cx43 hemichannels, detected within the non-junctional and soluble fraction of Cx43, have been demonstrated to participate in hyperexcitability and excitotoxicity (Almad et al., 2016, 2022; Guo et al., 2022; Rovegno et al., 2015). Furthermore, how local and spatially limited dysfunction of Cx43 contributes to the progression of brain pathology had not yet been addressed. In this study, we expanded our analysis of Cx43 expression and function following mild TBI. We found that Cx43 is upregulated at the protein level, yet Cx43 expression was highly heterogeneous from astrocyte to astrocyte within the cortical gray matter. Increased levels of Cx43 protein represented increased Cx43 hemichannel and/or cytoplasmic pools of Cx43, while GJ plaque density and size were largely unchanged. Inhibiting the phosphorylation of a key regulatory site of Cx43, serine 368, which maintains GJ opening while reducing hemichannel opening probability (Ek-Vitorin et al., 2006; Lampe et al., 2000), limited the seizure susceptibility of mice in both the presence and absence of mild TBI.

### Heterogeneous Cx43 levels after mild TBI within the same brain and between mice

One clear finding of the study is the high variability of Cx43 protein expression changes across the cortical gray matter within one individual, as well as between individuals. While we observed an increase in the expression of Cx43 after mild TBI, immunohistochemistry staining revealed a heterogenous expression pattern of the protein: some cortical areas had elevated Cx43, but others had a decrease or absence of it, highlighting the challenge of interpreting assays that rely on bulk assessments of Cx43 expression alone. This heterogeneity is at least in part caused by differences in the microenvironment. Previous studies in our laboratory showed that blood-brain barrier leakage induced an atypical astrocyte response characterized by significant reduction in astrocyte proteins including Cx43 that is not observed in other areas (George et al., 2021; Shandra et al., 2019). The increase of Cx43 in other areas could either compensate for Cx43 loss in atypical astrocyte areas and/or factors upstream of Cx43 phosphorylation may be responsible for the increase in Cx43 protein. To our knowledge, such heterogenous expression of Cx43 has not been described in the brain previously, and thus, investigating how this distinct local regulation affects the maintenance of local brain function needs to be studied. Furthermore, these results demonstrate the need for reassessing the interpretation of previous studies that evaluate Cx43 levels exclusively by western blot. Previously, global decrease of Cx43 has been proposed as a potential therapeutic approach to ameliorate TBI outcomes (Ichkova et al., 2019). Yet, the cellular mechanism by which Cx43 contributes to seizure susceptibility and brain pathology (increase of hemichannels, closing or loss of gap junctions, or functions of cytoplasmic Cx43) remains to be resolved.

We also found that the Cx43 global response varies between animals. Experiments that analyzed Cx43 expression levels, localization, or function all presented with high variability among individuals reminiscent of the highly variable nature of human TBI (Maas et al., 2022). Whether this variability is due to the variability of the injury itself or whether it is due to other factors regulating Cx43 is not yet clear.

### Cx43 increase after mild TBI is attributed to the non-junctional functions of Cx43

The role of Cx43 in forming gap junctions in the brain and other organs has led the field to correlate an increase in Cx43 protein expression with an increase in GJs. Yet, Cx43 has other functions unrelated to the GJ coupling of astrocytes. In this study, we differentiated between Cx43 GJs and non-junctional Cx43 using fractionation of Cx43 protein based upon solubility. We also used super-resolution imaging to identify Cx43 plaques, which are structures encompassing arrays of GJ channels (McCutcheon 2020). GJs are surprisingly dynamic structures, and alterations to plaque size occur rapidly in other systems during stress due, in part, to altered Cx43 phosphorylation (Smyth et al., 2014). Our experiments revealed that junctional Cx43 did not change in size or density of plaques. Instead, biochemical fractionation demonstrated increased ‘soluble’ Cx43 suggesting that mild TBI induces increased levels of hemichannels and cytoplasmic Cx43.

There are conflicting reports regarding the relevance of hemichannels in pathology (Nielsen et al., 2017). While some reports assign important roles to astrocytic Cx43 hemichannels, others question their physiological relevance. Despite such disagreement within the field, recent work has shown that Cx43 hemichannels contribute to increased excitotoxicity and pathology in amyotrophic lateral sclerosis (ALS) (Almad et al., 2022) and temporal lobe epilepsy (Guo et al., 2022). Yet, the field is just starting to understand the role of Cx43 hemichannels in other disorders.

In this study, we asked if the increase in soluble Cx43 was correlated with an increase in hemichannel function, taking advantage of the proportional relationship between EtBr uptake and the number of active hemichannels on the cell surface (Contreras et al., 2003; Hansen et al., 2014). Mild TBI increased CBX-dependent EtBr uptake into astrocytes suggesting increased activity of large pore (hemi)channels (Hansen et al., 2014). This uptake was not pannexin-dependent, as the pannexin 1 inhibitor spironolactone did not interfere with EtBr uptake. Using a Cx43-specific hemichannel inhibitor, EtBr uptake was reduced in 6 of 8 mice. This is in alignment with the above-discussed animal-to-animal variability that we observed across many outcome measures or could be due to a lack in diffusion efficiency of the TAT-Gap19 peptide in the 300 µm acute brain slices. Whether increased Cx43 hemichannel activity results in a release of excitotoxic compounds as reported in other disorders needs to be explored in the specific context of mild TBI.

### Cx43^S368^ phosphorylation contributes to seizure susceptibility after mild TBI

Cx43 function and location are tightly regulated by post-translational modifications. One of the key regulatory sites within the Cx43 C-terminus is serine 368, which upon phosphorylation via PKC effects GJ channel closure and precedes internalization following additional phosphorylation events and ubiquitination (Calhoun et al., 2020; Ek-Vitorin et al., 2006; Lampe et al., 2000; Sirnes et al., 2009). Previous studies assessed the levels of pCx43^S368^ in different models of TBI. In a study using lateral fluid percussion TBI, the authors found an increase in pCx43^S368^ in the hippocampus, which they correlated to an increase in hemichannel-released exosomes identified by labeling with CD81 and CD63 (W. Chen et al., 2018). Another study using controlled cortical impact TBI found an increase in Cx43 expression but no difference in pCx43^S368^ in the subventricular zone or the hippocampus (Greer et al., 2017). Finally, in cortical tissue of TBI patients increased pCx43^S368^ levels were positively correlated with injury severity and long-term TBI outcomes, including post-traumatic epilepsy (B. Chen et al., 2017). Human and experimental temporal lobe epilepsy samples also showed an increase in pCx43^S368^ (Deshpande et al., 2017). This correlative data is in alignment with our findings of increased pCx43^Ser368^ in the cortex after mild TBI.

To establish whether pCx43^Ser368^ is causally related to seizure susceptibility after mild TBI, we used mice with a serine to alanine substitution at Cx43^S368^, which interferes with phosphorylation of Cx43 at this residue. This conferred substantial protection from the development of seizures in both injured and uninjured Cx43^S368A^ mice after subthreshold PTZ injections. While control mice that incurred a mild TBI have a significantly lower seizure threshold than uninjured mice, this increase in seizure susceptibility no longer occurred when Cx43^S368^ remained unphosphorylated.

While this data suggests a key role for Cx43^S368^ in mediating pathological changes after TBI, one confounding factor in Cx43^S368A^ mice are substantially lower total Cx43 levels. Future studies need to resolve whether protection from seizures is due to reduced or increased gap junction function and/or attributable to other Cx43 functions, including hemichannels. Similarly, gap junction plaque densities were significantly lower even in uninjured Cx43^S368A^ mice, demonstrating that Cx43^S368^ does regulate GJs. Given that it has been suggested that GJs “feed” neuronal hyperexcitability in the context of pathology, reduced gap junction plaque densities in and of itself may be protective. Yet, the complete lack of Cx43 and Cx30 in astrocytes results in higher seizure susceptibility (Chever et al., 2016a; Deshpande et al., 2020) pointing to a different mechanism in Cx43^S368A^ mice, which have reduced seizure susceptibility despite the overall reduced Cx43 protein.

Importantly, the presence of a gap junction plaque does not reveal information about GJ conductivity. Previous work in cardiovascular infection/disease has demonstrated maintenance of gap junction coupling (Gy et al., 2011; Padget et al., 2024) while reduced hemichannel opening probability was reported (Hirschhäuser et al., 2021) in Cx43^S368A^ mice, providing evidence for either mechanism. While this data suggests that gap junctions in Cx43^S368A^ mice should be open at a higher probability and hemichannels should more likely be closed, this needs to be tested in brain.

In conclusion, this study demonstrates that mild TBI results in heterogeneous upregulation of Cx43 protein within the cortical gray matter, predominantly associated with non-junctional functions such as hemichannel activity rather than increased gap junction formation. Phosphorylation of Cx43^S368^, which reduces hemichannel opening while maintaining gap junction function, was found to decrease seizure susceptibility. Future studies need to delineate the specific contributions of gap junctions versus hemichannels in pCx43-mediated effects on seizure susceptibility in the brain.

## Acknowledgments

The authors want to acknowledge the contribution of Pratishtha Panigrani and Maame Boateng to this work, by organizing images and data. We thank Cynthia Wagner for overseeing EW’s work within the Applied Molecular Biology master’s program at the University of Maryland Baltimore County. The work in this manuscript was supported by the National Institutes of Health grants from the National Institute of Neurological Disorders and Stroke (R21NS107941 to SR and SL; R01NS121145 to SR), and the National Heart Lung and Blood Institute (R01HL159512 to JWS). Research carried out by CMB was supported by the National Institutes of Health Common Fund under Award Number U54CA272205. The content is solely the responsibility of the authors and does not necessarily represent the official views of the National Institutes of Health. OL was supported by the Cell and Molecular Biology Training Grant T32GM146611. RP was supported by the UAB Center for Community OutReach Development (CORD) Summer program. Research reported in this publication was conducted, in part, in the UAB High Resolution Imaging Facility.

## Table legends

**Suppl Table 1. Table with all results for PTZ experiments per animal.**

## Supplementary figures and table legends

**Supplementary Figure 1. Cx43^S368A^ mice genotype and phenotype validation for serine 368 to alanine substitution. A** Schematic of the portion of the *Gja1* gene (encoding Cx43) containing the point mutation and of the primers used in this study to confirm the genotype. **B** Table with conditions used to amplify the DNA fragment for genotyping. **C** Agarose gels showing the difference in band size between C57Bl/6 and Cx43^S368A^ mice. **D** Schematic of the NheI restriction enzyme recognition sequence within the part of *Gja1* containing the point mutation. **E** Agarose gels showing the different band sizes in Cx43^S368A^ and C57Bl/6 mice when that portion of the gene is amplified by PCR and after DNA was digested using NheI. **F** Western blot of cortical gray matter brain samples of C57Bl/6 and Cx43^S368A^ mice against pCx43^368^ and Cx43 proteins. Each lane represents a different animal. As an internal loading control, the membranes were probed for total protein. **G** Quantification of pCx43^S368^ presented as a ratio between pCx43^S368^ normalized to total protein and Cx43 normalized to total protein.

**Supplementary Figure 2. Cx43 IHC. A** Representative image of Cx43 gap junction plaques, Aldh1l1-tdTomato-positive (tdTom) astrocytes, and CD31-positive epithelial cells in the cortical gray matter. **B** Representative super-resolution image of vasculature in cortical tissue taken from the blue dotted box in Suppl. Fig. 2.A. Cx43 plaques colocalize with astrocyte processes and endfeet (arrows in the zoomed in box point to some examples for better orientation in the image). **C** A representative super-td-Tomato resolution image of larger vasculature in cortical gray matter tissue taken from the yellow dotted box in Suppl. Fig 2.A. Cx43 plaques colocalize with astrocyte processes and endfeet (arrows in the zoomed in box point to examples for better orientation in the image).

**Supplementary Figure 3. Uncropped western blots used to quantify Cx43 expression after mild TBI.** All western blots of cortical brain samples of C57Bl/6 animals used for quantification in Fig. 1 B. All samples were independent from each other (biological, not technical replicates) with each lane representing one mouse. Top panels and middle panels were imaged using infrared secondaries while the bottom image was developed using HRP-conjugated secondaries and chemiluminescence. Left panels show blots probed against Cx43 while the right panels show the same blot probed against GAPHD, which was used internal loading control for normalization. Excluded samples were indicated using an asterisk.

**Supplementary Figure 4. Uncropped western blots used to quantify Cx43 solubility.** All western blots used for quantifications in Fig. 2 B–D. Each lane represents either the soluble (sol), insoluble (ins), or total (tot) fraction of a sample. Red numbers indicate animal identity. Antibody used to probe the blot is indicated above the membrane image. Western blot membranes including total fractions (blot 3 and 4) were re-probed against GAPDH antibody and used for normalization.

**Supplementary Figure 5. The pannexin1-blocker spironolactone does not inhibit ethidium bromide uptake in astrocytes after mild TBI. A** Representative confocal image of control and spironolactone-treated astrocytes at 3 dpi. Green: S100β, magenta: EtBr. **B** Quantification of EtBr fluorescence intensity levels in control and spironolactone-treated slices at 3 dpi.

**Supplementary Figure 6. Uncropped western blot used for quantification of pCx43^S368^/Cx43.** All western blots of cortical gray matter brain samples of C57Bl/6 animals used for quantification in Fig. 4 B. Left blots were probed against pCx43^S368^ (top) and re-probed against GAPDH (bottom). Right blots were probed against Cx43 (top) and re-probed against GAPDH (bottom). Each lane represents one animal.

**Supplementary figure 7. Comparable results were obtained using rabbit Cx43 and panCx43 immunohistochemistry to assess Cx43 plaque density in C57Bl/6 and Cx43^S368A^ mice. A** Representative super-resolution images of Cx43 and panCx43 plaques at 3 dpi. **B** Quantification of Cx43 image intensity normalized to C57Bl/6 Sham (C57Bl/6 Sham mean = 100 AU ±10.15, n = 3; C57Bl/6 TBI mean = 100.8 AU ±7.91, n = 4; Cx43^S368A^ Sham mean = 73.86 AU ±3.699, n = 3; Cx43^S368A^ TBI mean = 77.23 AU ±4.612, n = 4; two-way ANOVA interaction p-value = 0.8601, treatment p-value = 0.7719, mouse strain p-value = 0.0055) and panCx43 image intensity normalized to C57Bl/6 Sham (C57Bl/6 Sham mean = 100 AU ±8.110, n = 3; C57Bl/6 TBI mean = 98.63.8 AU ±5.978, n = 3; Cx43^S368A^ Sham mean = 59.82 AU ±5.993, n = 3; Cx43^S368A^ TBI mean = 65.84 AU ±15.18, n = 3; two-way ANOVA injury p-value = 0.8143, genotype p-value = 0.0052, interaction between injury and genotype p-value = 0.7096). **C** Quantification of Cx43 plaque size normalized to C57Bl/6 Sham (C57Bl/6 Sham mean = 100 µm^2^ ±7.429, n = 3; C57Bl/6 TBI mean = 85.13 µm^2^ ±8.884, n = 4; Cx43^S368A^ Sham mean = 86.16 µm^2^ ±6.180, n = 3; Cx43^S368A^ TBI mean = 83.27 µm^2^ ±5.844, n = 4; two-way ANOVA injury p-value = 0.2625, genotype p-value = 0.3184, interaction between injury and genotype p-value = 0.4419) and panCx43 plaque size normalized to C57Bl/6 Sham (C57Bl/6 Sham mean = 100 µm^2^ ±14.19, n = 3; C57Bl/6 TBI mean = 105.0 µm^2^ ±9.591, n = 3; Cx43^S368A^ Sham mean = 79.81 µm^2^ ±3.774, n = 3; Cx43^S368A^ TBI mean = 93.39 µm^2^ ±5.681, n = 3; two-way ANOVA for injury: p-value = 0.3428, for genotype p-value = 0.1228, interaction between injury and genotype: p-value = 0.6542). **D** Quantification of Cx43 plaque density normalized to C57Bl/6 Sham (C57Bl/6 Sham mean = 100 plaque/µm^2^ ±4.590, n = 3; C57Bl/6 TBI mean = 84.17 plaque/µm ±11.84, n = 4; Cx43^S368A^ Sham mean = 47.42 plaque/µm^2^ ±6.763, n = 3; Cx43^S368A^ TBI mean = 41.99 plaque/µm ±4.437, n = 4; two-way ANOVA injury: p-value = 0.2254, genotype p-value = 0.0002, interaction between injury and genotype: p-value = 0.5412) and panCx43 plaque density normalized to C57Bl/6 Sham (C57Bl/6 Sham mean = 100 plaque/µm^2^ ±16.99, n = 3; C57Bl/6 TBI mean = 124.9 plaque/µm^2^ ±13.23, n = 3; Cx43^S368A^ Sham mean = 30.26 plaque/µm^2^ ±6.453, n = 3; Cx43^S368A^ TBI mean = 39.73 plaque/µm^2^ ±10.42, n = 3; two-way ANOVA injury: p-value = 0.6871, genotype: p-value = 0.0006, interaction between injury and genotype: p-value = 0.7206). In all graphs, Tukey’s multiple comparisons for group-to-group comparisons was reported as p-values.

**Supplementary figure 8. Cx30 gap junction plaques are unchanged in Cx43^S368A^ mice. A** Representative super-resolution images of Cx30 immunohistochemistry at 3 dpi. **B** Quantification of Cx30 plaque size 3 dpi (C57Bl/6 Sham mean = 0.08893 µm^2^ ± 0.01048, n = 3; C57Bl/6 TBI mean = 0.09672 µm^2^ ±0.01681, n = 4; Cx43^S368A^ Sham mean = 0.1615 µm^2^ ± 0.07716, n = 3; Cx43^S368A^ TBI mean = 0.2280 µm^2^ ± 0.1010, n = 3; two-way ANOVA test injury p-value = 0.5457, genotype p-value = 0.1193, Interaction between injury and genotype p-value = 0.6315; Tukey’s multiple comparison test used for group to group comparisons, p-values reported in figure). **C** Quantification of Cx30 plaque density 3 dpi (C57Bl/6 Sham mean = 0.4020 plaque/µm^2^ ±0.1358, n = 3; C57Bl/6 TBI mean = 0.6688 plaque/µm ±0.1774, n = 4; Cx43^S368A^ Sham mean = 0.7451 plaque/µm^2^ ±0.1641, n = 3; Cx43^S368A^ TBI mean = 0.8601 plaque/µm ±0.1024, n = 3; two-way ANOVA test injury p-value = 0.2254, genotype p-value = 0.0002, Interaction between injury and genotype p-value = 0.5412; Tukey’s multiple comparison test used for group to group comparisons, p-values reported in figure). **D** Representative western blot of cortical brain samples of C57BL/6 and Cx43^S368A^ mice against Cx30 and corresponding total protein. **E** Quantification of Cx30 protein expression normalized to total protein and C57BL/6 (C57Bl/6 mean = 100±2.814, n = 4; Cx43^S368A^ mean = 99.13±4.886, n = 4; unpaired t test, p = 0.8824). P-value of the statistically significant comparison shown in the graph.

