## Supplementary figures and images for "Astrocytic connexin43 phosphorylation contributes to seizure susceptibility after mild traumatic brain injury"

### Suppl Fig 7

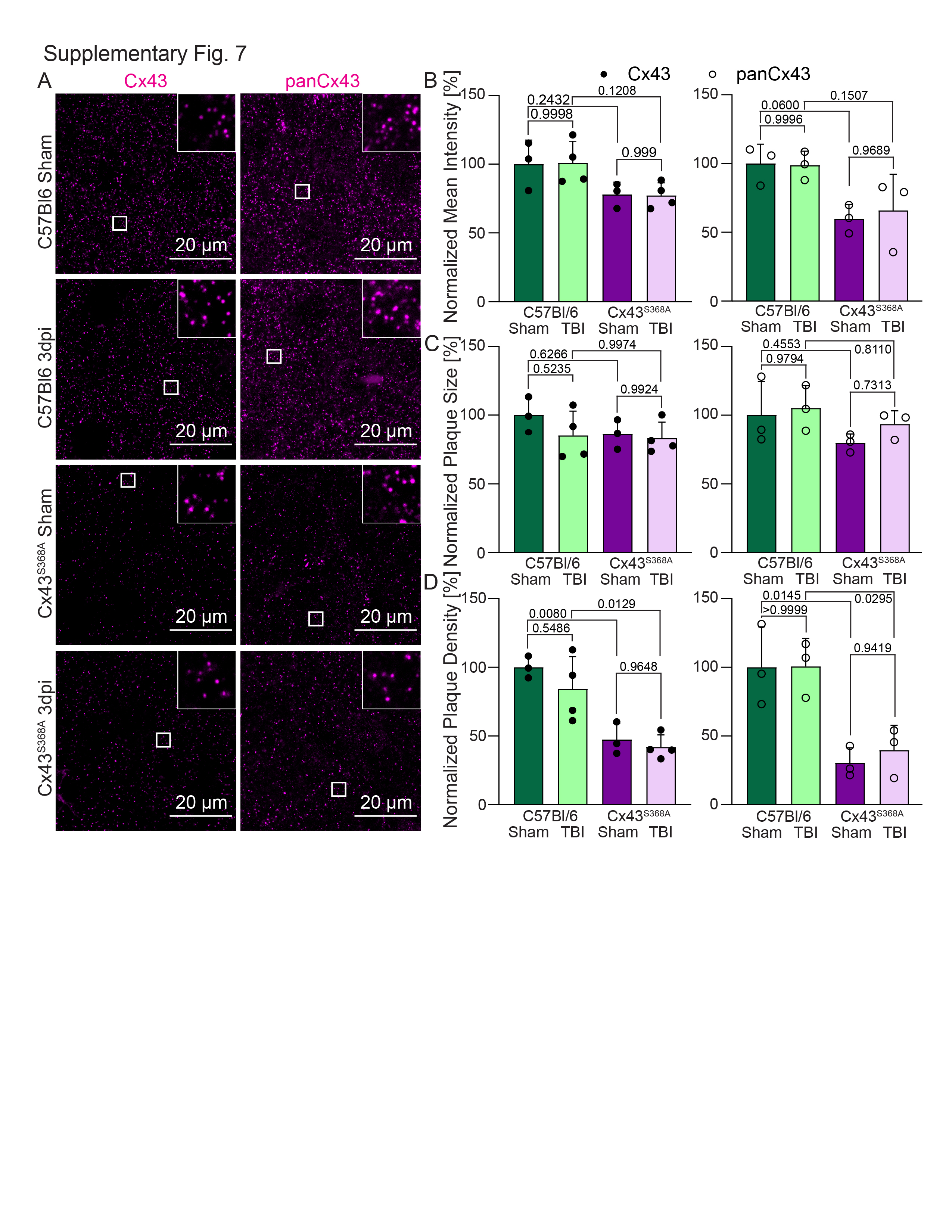

### Suppl Figure 1

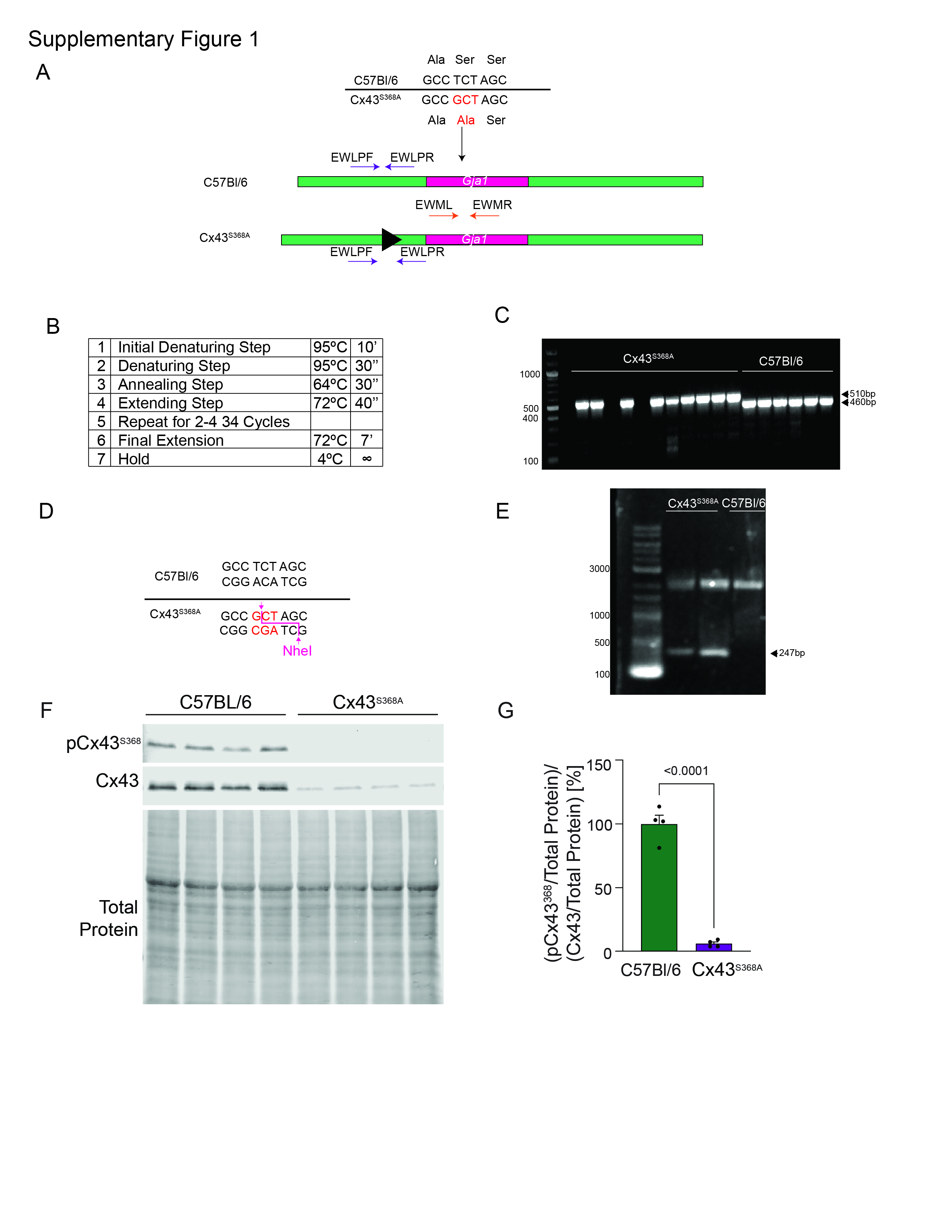

### Suppl Figure 2

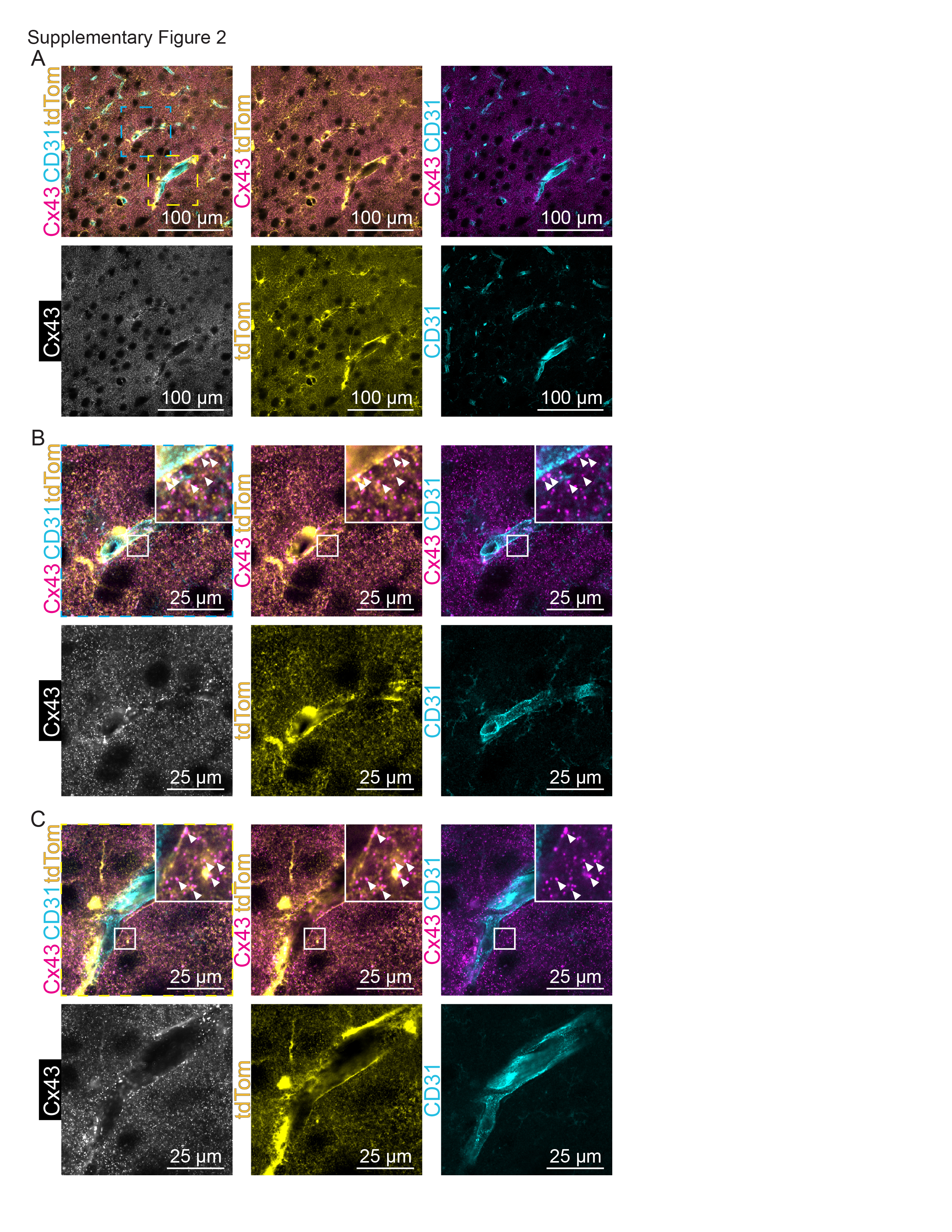

### Suppl Figure 3

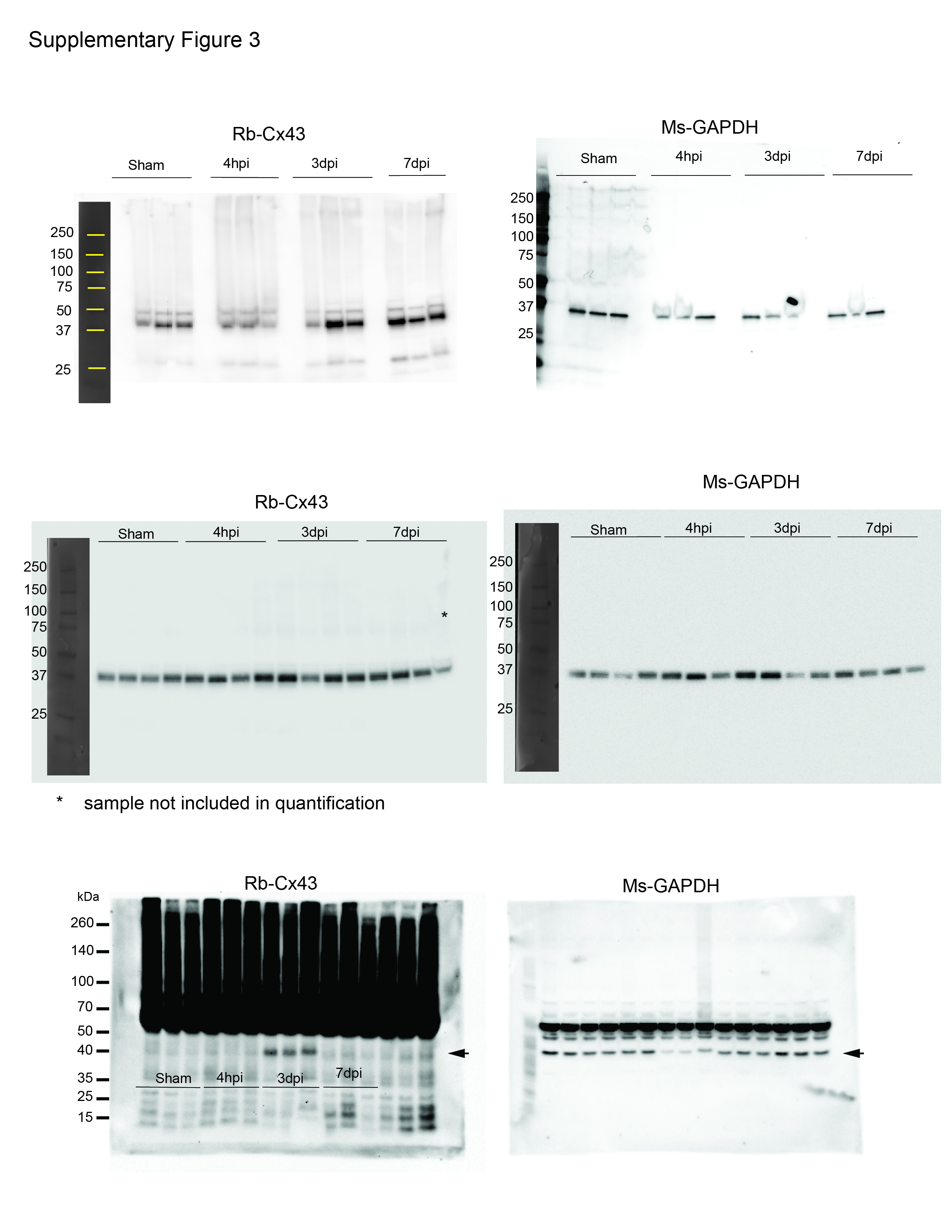

### Suppl Figure 4

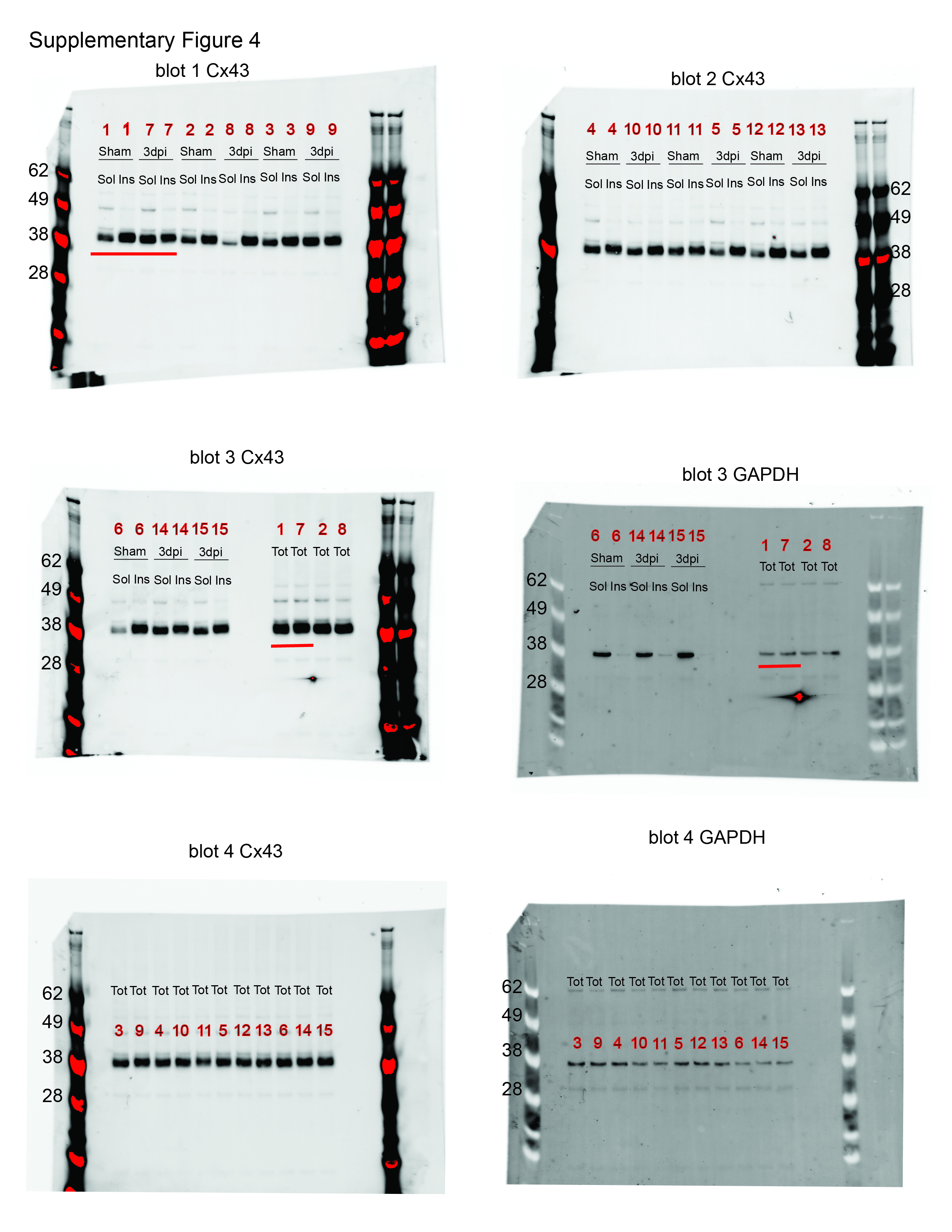

### Suppl Figure 5

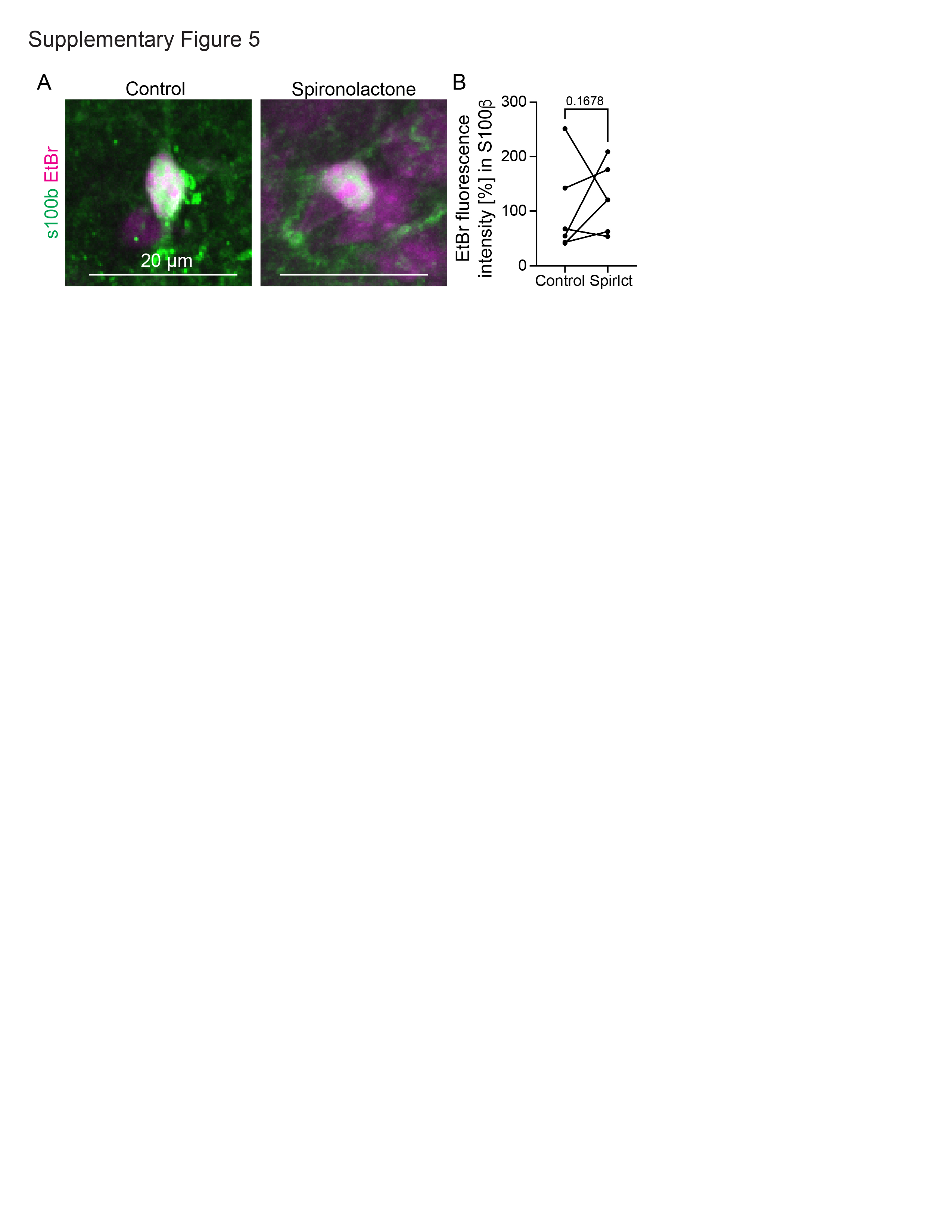

### Suppl Figure 6

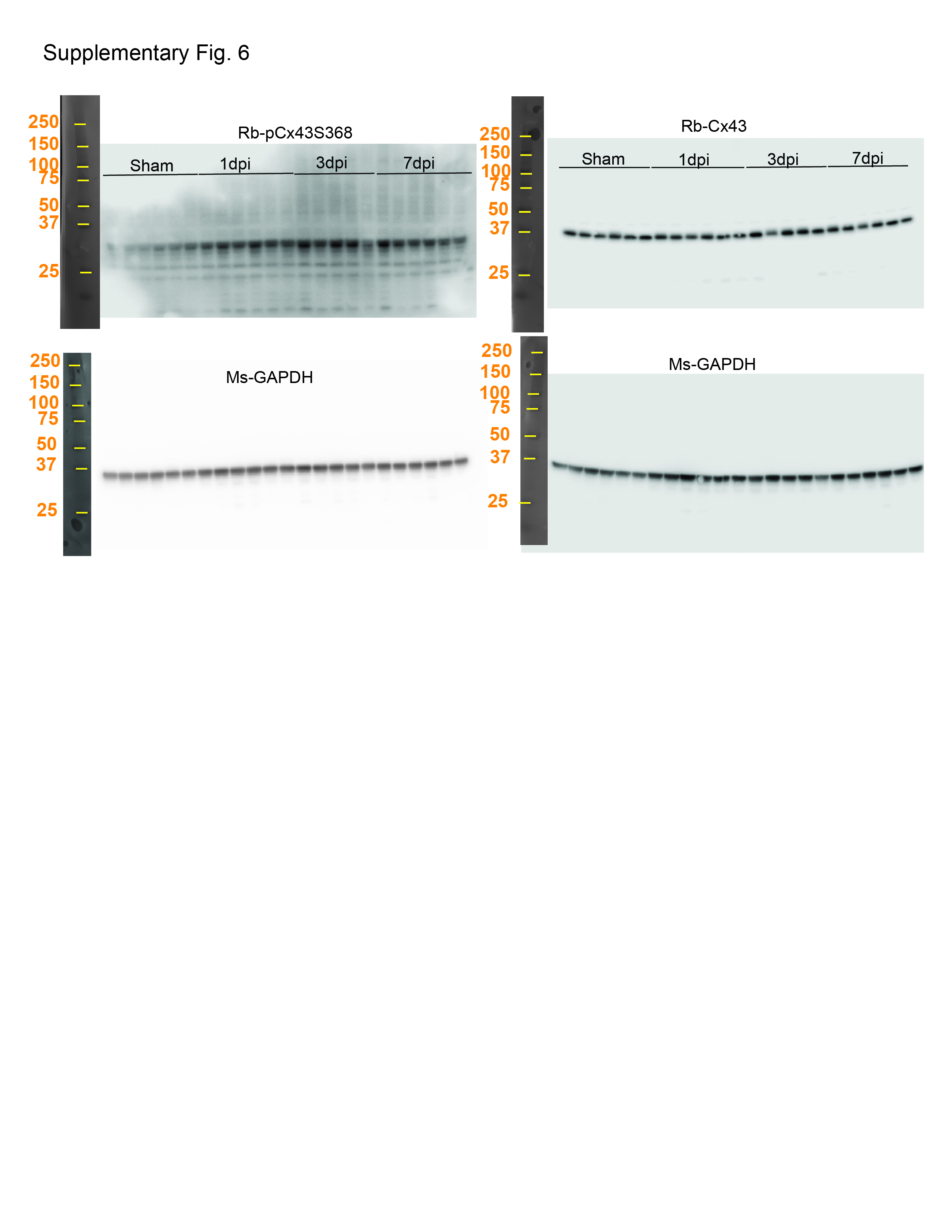

### Suppl Figure 8

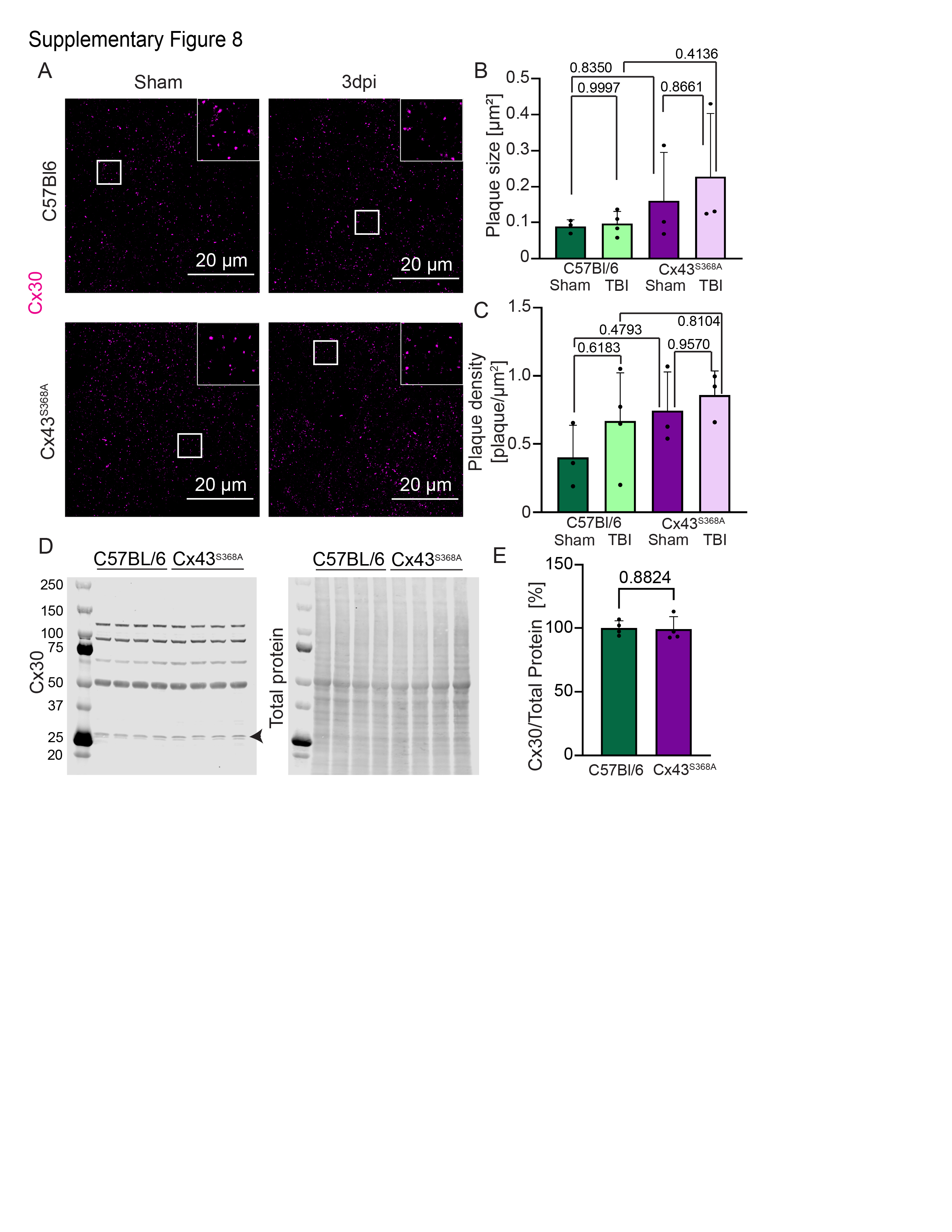
